# Sleep stage-specific effects of 0.75 Hz phase-synchronized rTMS and tACS on delta frequency activity during sleep

**DOI:** 10.1101/2025.05.09.653171

**Authors:** Kuri Takahashi, Min-Fang Kuo, Michael A. Nitsche

## Abstract

Slow oscillatory activity during non-rapid-eye-movement (NREM) sleep plays a crucial role in both physical health and cognitive functions. Enhancing slow oscillatory activity during sleep has the potential to benefit these domains, yet an optimal stimulation protocol has not been established. This study aimed to investigate whether repetitive transcranial magnetic stimulation (rTMS), synchronized with the trough phase of 0.75 Hz transcranial alternating current stimulation (tACS) can modulate EEG activity in the delta frequency range during sleep and enhance cognitive functions. Healthy adults participated in a within-subject, counterbalanced study design comparing real and sham stimulation conditions. Combined rTMS and tACS was applied over the bilateral prefrontal cortex before sleep. We evaluated (1) power spectral density and functional connectivity within the delta frequency range during resting state and sleep, (2) retention of declarative memory learned before sleep, and (3) sleep parameters including spindle activity, sleep stage ratios, sleep onset latency and sleep efficiency. The combined rTMS and tACS protocol significantly increased delta oscillatory activity during the N3 sleep stage compared to sham. Functional connectivity, as measured by global efficiency, was enhanced during the N2 sleep stage. However, the stimulation did not improve declarative memory retention, spindle activity or other sleep parameters. These findings demonstrate the potential of combined rTMS and tACS as a non-invasive method to enhance delta oscillatory activity during sleep. While the stimulation did not improve memory performance, its ability to modulate delta activity during sleep suggests potential clinical applications for addressing pathological alterations in slow wave activity during sleep.

**State of Significance:** Slow oscillatory activity during non-rapid eye movement (NREM) sleep is relevant for declarative memory consolidation and overall sleep quality. However, noninvasive brain-stimulation methods to selectively modulate these oscillations remain elusive. In this study, we show that pre-sleep application of repetitive transcranial magnetic stimulation (rTMS) synchronized to the trough of 0.75 Hz transcranial alternating current stimulation (tACS) significantly enhances delta oscillatory activity during N3 sleep and increases global functional connectivity during N2 sleep in healthy humans without sleep disturbances without altering sleep structure. Our findings indicate that rTMS synchronized to the trough of the tACS exerts sleep-stage-specific effects on delta oscillatory activity. This stimulation protocol represents a promising approach for restoring pathological slow-wave activity during sleep in clinical populations.

## Introduction

More than simply being relevant for rest, sleep plays a critical role in maintaining psychological health, supporting the immune system, and enhancing cognitive processes, particularly memory consolidation ^1–3^. Memory consolidation refers to the process of transforming newly acquired information into long-term storage. Sleep is divided into five distinct stages: wakefulness, N1, N2, N3, and REM sleep ^4^. Some of these stages are relevant for consolidation of specific memory types such as declarative, procedural, and emotional memory ^5^.

Among these stages, N3 sleep, also known as slow-wave sleep (SWS), is particularly relevant for the consolidation of hippocampus-dependent declarative memories ^6^. SWS is characterized by high-amplitude, low-frequency delta oscillatory activity (0.5-4 Hz), originating primarily from thalamic neurons ^4,7,8^. During SWS, reactivation of newly encoded memories within the hippocampus and memory transfer from the hippocampus to the neocortex takes place ^9,10^. In accordance, a positive correlation between SWS and declarative memory performance improvement, and an association between delta oscillations during SWS and memory performance improvement have been demonstrated ^11–13^. Beyond the contribution of delta oscillations, sleep spindles, burstlike oscillatory EEG activity originating from the thalamus, also play an important role in memory consolidation ^14,15^. By inducing synaptic plasticity, which strengthens neural connections and provides a foundation for memory stabilization, spindles aligned with slow oscillations facilitate the transfer of hippocampal memory traces to the neocortex ^16,17^. At the cognitive level, this enhances memory, making it resistant to interference, and enhances performance without additional practice ^3^.

Moreover, the significance of delta oscillatory activity during SWS is not limited to the consolidation of memory, but also crucial for maintaining overall sleep quality. Disruptions of delta activity have been linked to sleep disorders, such as insomnia ^18,19^. While memory consolidation in healthy individuals benefits from SWS, insomnia patients with reduced SWS show respective deficits ^20,21^. This underlines the importance of delta oscillatory activity for both, health and cognitive functions.

Given the role of delta oscillations in overall sleep quality and sleep-dependent memory consolidation, non-invasive brain stimulation (NIBS) has emerged as a promising intervention to enhance delta activity during sleep ^22,23^. Marshall et al. (2004, 2006) demonstrated that 0.75 Hz slow-oscillatory transcranial direct current stimulation (so-tDCS) - which combined an oscillatory sinusoidal waveform with anodal tDCS in that study – applied during SWS increased slow-wave EEG activity in the frontal cortex. This enhancement was accompanied by increased spindle activity and improved declarative memory performance.

The DC offset in so-tDCS may lead to sustained alterations of cortical excitability by inducing synaptic plasticity including long-term potentiation (LTP) – like changes ^24^. The DC offset in so-tDCS may have led to a sustained alteration of the resting membrane potential and, consequently, prolonged cortical excitability alteration ^25,26^. Given this characteristic, so-tDCS may also induce long-term potentiation (LTP)-like neuroplasticity ^24^, a mechanism underlying memory formation in both the hippocampus and cerebral cortex ^27^. Thus, it remains unclear whether the observed improvements of declarative memory during sleep observed after so-tDCS are driven purely by increased delta and spindle activity or by other mechanisms such as neuroplasticity.

A recent study from our group has demonstrated that repetitive transcranial magnetic stimulation (rTMS) synchronized to the trough of the transcranial alternating current stimulation (tACS) waveform led to significant and sustained enhancement of delta oscillatory activity and functional connectivity during wakefulness ^28^. The efficacy of this combined approach in other frequency ranges such as theta and alpha frequencies was also supported by previous studies ^29,30^.

Unlike tDCS, tACS modulates endogeneous neuronal activity entraining it to the targeted frequency band. However, the efficacy of tACS relies on the prior endogenous neuronal activity at the targeted frequency band ^31,32^. rTMS, on the other hand, induces frequency-specific modulation by repeatedly delivering pulses at a specific frequency, though its effects are typically short-lasting ^33^. Thus, it has been assumed that, by stabilizing rTMS-induced oscillatory activity, tACS can prolong and strengthen rTMS effects and cancel out the limitations of each technique. Under this assumption, previous studies have demonstrated that combining these stimulation techniques enhances the modulation of the targeted frequency, with improvements in cognitive performance and functional connectivity sustained for up to one hour compared to using either technique alone ^31,32^.

Based on these findings, we hypothesized that combined delta-frequency rTMS and tACS, previously shown to induce and stabilize delta oscillatory activity during wakefulness would similarly lead to changes in delta activity during sleep, given its potential to stabilize delta oscillations over extended periods. In addition, we hypothesized that this increased delta activity would enhance declarative memory consolidation, as suggested by a previous study^23^.

In the current study, we applied either delta-frequency rTMS and tACS or sham rTMS and sham tACS to healthy adults before sleep. We compared changes in delta power and functional connectivity in the delta frequency band during sleep between real and sham stimulation conditions. Additionally, we measured delta power changes during the resting state before sleep to enable a comparison with our previous study that applied stimulation during the resting state and reported enhanced delta power ^28^ We also measured resting-state EEG after sleep to explore the duration of the stimulation after-effects. Moreover, we aimed to replicate the findings of Marshall et al. (2006) by investigating whether enhancing delta activity leads to changes in spindle activity and improves declarative memory consolidation.

## Materials and methods

### Participants

Sixteen healthy young adult participants were recruited (8 males, mean age 25.06, SD 3.03). Exclusion criteria were diagnosis of insomnia disorder, non-German native speaker, tobacco smokers, pregnancy, history of neurological or psychiatric disorders, including epilepsy or head trauma, intake of CNS-acting medication, and having metal implants. Prior to the beginning of the experimental sessions, all participants were screened by a medical professional and confirmed to have normal sleep. All participants provided written informed consent and were financially compensated for participation. The experiment conformed to the principles of the Declaration of Helsinki and was approved by the Institutional Review Board of the Leibniz Research Centre for Working Environment and Human Factors (Dortmund, Germany). The datasets generated in the current study are available from the corresponding author upon reasonable request.

### Stimulation

We used the combined rTMS and tACS protocol which was found to be most effective to induce oscillatory delta activity in the previous study of our group ^28^. The duration of the intervention was chosen in a manner compliant with technical and safety limits ^34^. We employed a custom-made circuit to align transcranial magnetic stimulation (TMS) pulses within the trough phase of the tACS waveform, which was described in detail in previous studies ^28–30^. Figure 1 shows an overview of the circuit, and depicts the waveform and the timing of the stimulation conditions. Biphasic TMS pulses were delivered using two figure-of-eight shaped coils (diameter of each winding 70 mm, PMD coil type; Mag and More GmbH, Munich, Germany). For the sham condition, two separate sham coils were used (double 70 mm pCool-SHAM coil; Mag and More GmbH, Munich, Germany) which elicited audible clicks at the same decibel intensity as the active/real stimulation coil, but without inducing electromagnetic pulses. In order to deliver TMS pulses through two coils with no latency between pulses, the TMS devices were connected in series with TTL signals triggering the TMS. The two pulses were verified to have zero latency via measurements obtained by an oscilloscope. With regard to tACS, we used a Starstim 8-channel constant-current, battery-powered electric stimulator (Neuroelectrics, Barcelona, Spain). Stimulation electrodes were circular (2 cm radius, 12.57 cm^2^ area) and made of carbonated rubber, with the connector located at the center of the electrode. Conductive Ten20 paste (Weaver and Company, USA) was applied to the bottom of the electrodes. Care was taken to ensure that the electrodes had full contact with the scalp, with hair moved to the side. Stimulation electrodes were positioned over the F3, F4, TP9 and TP10 EEG electrode positions (10-20 International System).

**Figure 1.**
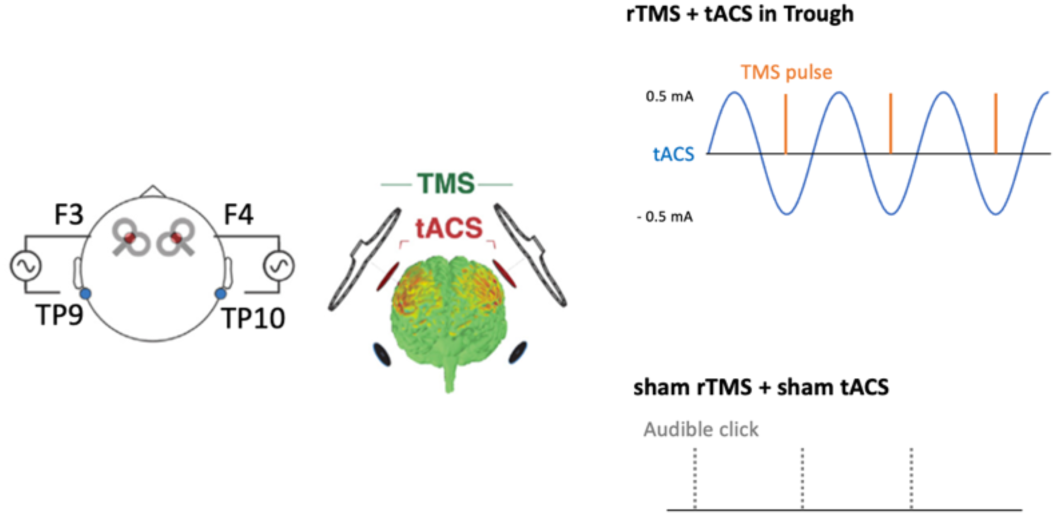
The rTMS and tACS stimulation montage and timing of stimulation application in real stimulation and sham stimulation protocols are shown. Orange bars indicate TMS pulses, gray dotted bars indicate audible clicks of sham coils, and blue waveforms represent a single tACS channel with a peak-to-peak amplitude of 1 mA. The montage consisted of two TMS coils and four tACS electrodes. TMS coils were positioned at an angle of 45° to the midline, aligned with each target tACS electrode during stimulation. Two tACS electrodes applied sinusoidal current in the same phase over F3 and F4 EEG positions, while the remaining two electrodes applied currents in the opposite phase over the left and right mastoid positions (TP9 and TP10). In the real stimulation condition, rTMS was delivered at the negative peak of the tACS waveform. In the sham condition, only audible clicks from sham TMS coils were applied (The figure was adapted from Hosseinian et al. (2021a,b) and modified with permission).

The multichannel Starstim device allows external programming of customized stimulation waveforms at fixed latencies and current amplitudes through the eight available channels. As such, two channels were used to program pulses at precise timepoints for delivering rTMS pulses in predefined relation to four other channels that were used to deliver sinusoidal alternating currents to the head.

For triggering TMS, pulses (0.1 mA peak, 0.5% duty cycle) were delivered from the electric stimulator to a separate circuit containing an optocoupler (developed in-house) in order to shield the stimulation from the main power. Current-to-voltage conversion resulted in a 5 V signal which was used as a TTL signal to communicate with the TRIGGER-IN of the TMS machine. Latencies of the waveform generated by the electric stimulator and the TMS stimulator were measured and verified to be aligned (+/- 1 ms resolution) via an oscilloscope. As mentioned above, tACS included four electrodes, with two channels delivering alternating current in the same phase to the left and right prefrontal cortex, and antiphase to the respective ipsilateral mastoid region. Stimulation electrodes were positioned at the F3, F4, TP9 and TP10 EEG positions. Two TMS coils were positioned directly over each tACS electrode at F3 and F4, and received signals from the tACS device through the circuit which triggered pulses at precise timepoints relative to the tACS sinusoid. Both, TMS and tACS were delivered with the frequency of 0.75 Hz for 30 min continuously. rTMS was applied at 80% of each individual’s active motor threshold, while tACS was applied at 1 mA peak-to-peak intensity. Sham tACS included a brief ramp-up over 15 seconds, immediately followed by a ramp-down over 15 seconds at the beginning of the stimulation to mimic the cutaneous sensation of real tACS. The TMS coils were held and maintained by the coil holders over the F3 and F4 stimulation sites. Electrode cables were only connected during the course of stimulation, and then detached for the remainder of the recording.

### Resting-state EEG

*Recording:* Resting-state EEG was recorded with a NeurOne Tesla EEG amplifier (Bittium, NeurOne, Bittium Corporation, Finland). 62 Ag/AgCI EEG electrodes were mounted on the head with a cap (EASYCAP GmbH, Herrsching, Germany) and a high-viscosity electrolyte gel (SuperVisc, Easycap, Herrsching, Germany) was used to establish a connection with the head. The reference electrode was positioned on the left mastoid, and the ground electrode was placed at the CPz position. During recording, digital TTL triggers transmitted to the EEG system by a presentation PC were used to mark the start and end of all experimental conditions.

#### Offline preprocessing and analysis

Offline preprocessing of resting-state EEG was performed using MNE-Python ^35^. The raw data were downsampled to 512 Hz. Electrodes contaminated by noise were identified using the Random Sample Consensus (RANSAC) algorithm implemented in the Python package, *autoreject* ^36^. The data were then re-referenced to the average of the EEG channels, excluding artifactual electrodes, as well as the F3, F4, TP9 and TP10 electrodes, which were located in areas covered by the tACS electrodes. A high-pass filter at 0.1 Hz and a low-pass filter at 45 Hz were applied offline. To further reject artifacts, an independent component analysis (ICA) using the *fastica* algorithm was applied ^37^. Components associated with eye blinks were automatically detected and removed based on Pearson correlation between each independent component and the EOG channel. This detection method employed an adaptive z-scoring technique, identifying components with z-scores greater than or equal to 3 as blink-related. These components were removed iteratively until no further blink-related artifacts were detected. After cleaning the data, artifactual electrodes were interpolated per subject using the cleaned data.

To examine changes in delta-band oscillatory activity, the power spectral density (PSD) of predefined frontal, central and temporal channels (spanning the region of stimulation; F5, F6, FC3, FC4, FC5, FC6, C5, C6, C3, C4, F7, F8, FT7, FT8, T7, T8, TP7, TP8, CP5, CP6, CP3, CP4) was computed at 0.75 Hz as well as across the entire delta band (0.5-3.9 Hz). PSD calculations were performed using the multitaper method ^38^ with the default window half-bandwidth of 4 Hz. Outliers in raw power values were identified as data points falling below the first quartile (Q1, −1.5 times of the interquartile range (IQR)) or above the third quartile (Q3, + 1.5 times of the IQR), and were excluded from further analysis.

### Polysomnography

#### Recording

Polysomnography was recorded using a NeurOne Tesla EEG amplifier (Bittium, NeurOne, Bittium Corporation, Finland). Neural activity was measured using 62 Ag/AgCI EEG electrodes, two EOG electrodes to track vertical and horizontal eye movements, and one EMG electrode under the chin. The sampling frequency was set at 2000 Hz, and the 62 electrodes were mounted on the head with a cap (EASYCAP GmbH, Herrsching, Germany). The reference electrode was positioned on the left mastoid, and the ground electrode was placed at the AFz position. To establish a connection with the head, we used a high-viscosity electrolyte gel (SuperVisc, Easycap, Herrsching, Germany). To ensure the cap remained in its original position even if participants moved, the cap was fixed using gauze. EEG impedance maintained at values less than 10 kOhms throughout the recording.

#### Offline preprocessing and analysis

The raw data were first downsampled to 512 Hz, and polysomnography signals were segmented into 30-second epochs. For further preprocessing and analyses, the Python package *Yet Another Spindle Algorithm* (YASA) ^39^, specifically designed for analyzing polysomnographic sleep recordings, was used. Artifactual epochs were identified and excluded from further analysis using YASA’s artifact detection function to ensure that only robust epochs were retained for analysis. Artifact detection was based on the standard deviations of each epoch. A moving window of three seconds was applied to calculate z-scores across channels, and epochs with z-scores exceeding a threshold of 3 were flagged as artifacts. Sleep stages were scored using YASA’s sleep staging function, which applies frontal, central and occipital derivations (F1, F2, C3, C4, O1, and O2) based on the American Academy of Sleep Medicine (AASM) guideline ^4^. The sleep stage scoring algorithm in YASA employs a machine learning-based classification model that extracts features from the data and labels each epoch based on characteristic patterns associated with the respective sleep stages ^39^. Using the scored polysomnographic data, sleep statistics including total sleep time, time spent in each sleep stage, sleep efficiency and sleep onset latency were computed for each participant using YASA’s sleep statistics function.

To analyze delta-band oscillatory activity during sleep, the PSD of predefined frontal, central and temporal channels (spanning the region of stimulation; F5, F6, FC3, FC4, FC5, FC6, C5, C6, C3, C4, F7, F8, FT7, FT8, T7, T8, TP7, TP8, CP5, CP6, CP3, CP4) was computed at 0.75 Hz and across the entire delta band (0.5-3.9 Hz). PSD calculations were performed for all epochs using the multitaper method ^38^, with the default window half-bandwidth of 4 Hz. Outliers in raw power values were identified as data points falling below the first quartile (Q1, −1.5 times of the interquartile range (IQ R)) or above the third quartile (Q3, + 1.5 times of the IQR), and were excluded from further analysis.

Additionally, sleep spindles were analysed in frontal (F5, F1, Fz, F2, F6) and parietal (P5, P3, P1, Pz, P2, P4, P6) channels using the sleep spindle detection algorithm in the YASA package, *spindles_detect*, based on the sleep-spindle detection method suggested by Lacourse et al. (2019) ^40^ for N2 and N3 sleep stages. Spindle detection was performed separately for slow (9-11.9 Hz) and fast sleep spindles (12-15 Hz) ^41,42^.

### Whole brain functional connectivity analyses

Whole brain functional connectivity analyses were conducted following the methods described in our previous study ^28^. Specifically, connectivity information during resting-state and sleep was captured between all pairwise combinations of EEG channels using the weighted phase lag index (wPLI), as implemented in the Python package *MNE-Connectivity*^35^. EEG data were first segmented into 10-second epochs, and the connectivity analysis was performed on the epoched data. For the polysomnography data, the connectivity analysis was performed iteratively for each sleep stage.

The wPLI is defined as:

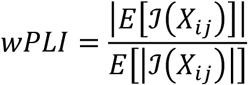

where *X_ij_* represents the cross-spectral density (CSD) between two signals *S_i_*(*t*) from channel *j* and *S_j_*(*t*, from channel *j*. ℐ refers to the imaginary component, and *E*[] is the expectation ^43^.In these equations, *X_ij_* is the cross-spectral density (CSD) between two signals *S_i_*(*t*) from channel *j* and *S_j_*(*t*, from channel *j*. ℐ refers to the imaginary component, and *E*[] is the expectation ^43^.

The CSD *X_ij_* of channel *j* and *j* is defined as:

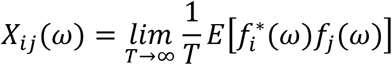

where *f_j_*(ω) is the finite Fourier transform of signal *S_j_*(*t*, at frequency ω. *f_i_*^∗^(ω) is the complex conjugate of *f_i_*(ω) at frequency ω.

At the whole-brain level, we adopted global efficiency *E*_global_ as a measure of global information integration. Resting state EEG was used to compute global efficiency, a metric that quantifies the capacity of the brain to integrate information. A previous study has shown that this metric accurately measures information integration capacity and is associated with improvements of working memory performance ^30^.

*E*_global_ is defined as:

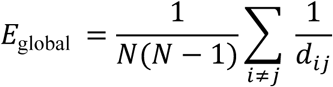

where *N* is the number of nodes in the network *G*. *d*_j,k_ is the shortest path between nodes *i* and *j*. We used wPLIs as weights of edges to construct the graph without thresholding the weights. Global efficiency is a scaled measure ranging from 0-1, with a value of 1 indicating maximum global efficiency of a network. This measure was computed by the network graph constructed by the wPLIs using the *Brain Connectivity Toolbox* ^44^.

### Paired associate learning task

We employed a paired associate learning (PAL) task, which was also used in previous studies^22,23^. The experiment was programmed and presented using the PsychoPy software package ^45^. The procedure of the PAL task was identical to that described in previous studies ^22,23^.

A different word list was used for each of the two experimental sessions. Each list consisted of 46 pairs of German nouns. The two word lists were designed as parallel versions and were assinged in a counterbalanced manner across participants to control for potential order effects. Additionally, four dummy word pairs were included at the beginning and end of each list to buffer primacy and recency effects. The sequence of word-pair presentations was randomized in each trial.

In the learning condition before sleep, each word pair was displayed on a monitor with a presentation rate of 0.20 seconds and an interstimulus interval of 100 milliseconds. The presentation of word pairs was followed by a cued recall task. During the cued recall, only the 46 cue words were presented on the screen in a different order from the initial presentation.

Participants were given unlimited time to recall the paired words. If participants failed to achieve a minimum of 60% correct responses, the word pairs were presented again in a newly randomized order, and the cued recall task was repeated.

In the recall condition after sleep, the 46 cue words were presented in a newly randomized order, and participants were given unlimited time to recall the corresponding paired words.

### Experimental design and procedure

The study was designed to assess whether rTMS trough-synchronized to tACS affects delta activity during sleep and memory performance compared to sham stimulation. The two experimental conditions were: 1) rTMS at the trough of the tACS wave; 2) sham rTMS combined with sham tACS. The study employed a within-subject, counterbalanced, single-blinded design.

Figure 2 depicts the overall experimental procedure. Each participant attended three sessions in total. The first session was designed as an adaptation to mitigate the first-night effect ^46^, followed by two experimental sessions with real and sham conditions, separated by at least seven days to prevent carry-over effects.

**Figure 2.**
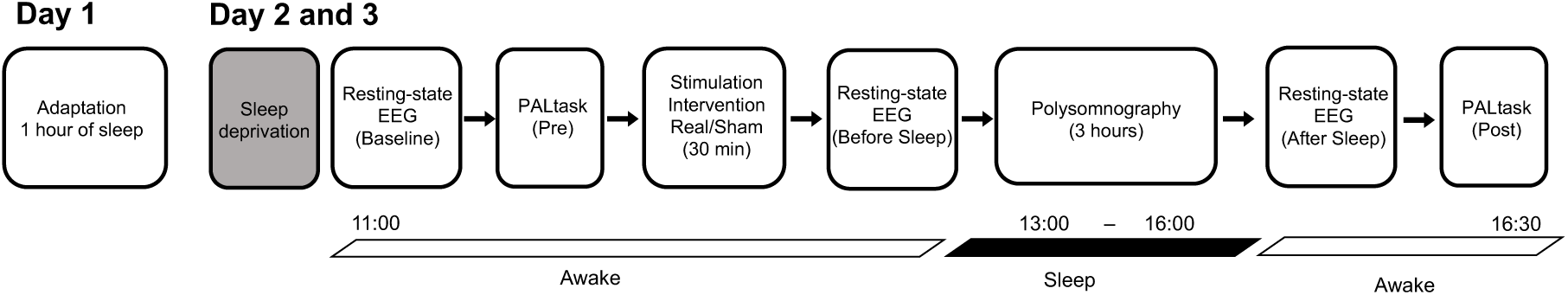
Experimental design. The study consisted of three sessions: an adaptation session to mitigate the first-night effect, followed by two experimental sessions (real and sham stimulation conditions). Participants underwent sleep deprivation before arriving at the sleep laboratory. Baseline resting-state EEG was recorded first. After completing the learning phase of the PAL task, participants received stimulation. rTMS was applied at 80% intensity of the individual active motor threshold (AMT), while tACS was applied continuously for 30 min at 1mA peak-to-peak intensity between the pairs of tACS electrodes. Resting-state EEG was recorded before and after 3 hours of sleep. Following sleep, participants completed the recall phase of the PAL task.

During the adaptation session, participants were asked to sleep in a sound-shielded sleep laboratory with an EEG cap. To induce sleepiness and ensure sufficient sleep during the daytime laboratory session, participants were instructed to stay awake from 9 pm the night before until their arrival at the laboratory. During this period of sleep deprivation, participants wore an actigraph (Motion Watch 8®, Cambridge Neurotechnology, Cambridge, UK) to confirm wakefulness. Participants did not consume caffeine or alcohol the night before the experimental session to prevent interference with sleep quality.

Each session began at 9 am. At the start of each session, a 62-channel EEG cap was prepared on the participant’s head according to the 10-10 convention. Baseline EEG recordings were conducted in eyes open (EO) and eyes closed (EC) conditions for two minutes each while participants were seated comfortably in a chair. Participants were then asked to learn the word pairs of the PAL task. Following this, the motor cortical hotspot was identified over the left primary motor cortex using TMS, and the active motor threshold (AMT) was determined by applying single TMS pulses to elicit motor-evoked potentials (MEP) from the right abductor digiti minimi (ADM) muscle. The lowest TMS pulse intensity required to elicit an MEP response of ∼200–300 μV during moderate tonic contraction of the right ADM muscle (∼20% of the maximum muscle strength) in at least three out of six consecutive trials was defined as AMT ^47^. For each individual, 80% of the stimulation intensity required to produce the AMT was used as rTMS intensity for the main experimental sessions. Following the acquisition of AMT, stimulation of one of the predefined rTMS/tACS combinations was delivered for 30 min. During stimulation, neuronavigation was performed with the Localite TMS Nagivator software (LOCALITE Biomedical Visualization Systems GmbH, Sankt Augustin, Germany) to monitor stability of the coil position. Following stimulation, resting-state EEG recordings were conducted in EO and EC conditions while participants lay on the bed. After resting-state EEG recordings, the lights were turned off, and participants were instructed to sleep. Total sleep duration was approximately three hours for all participants (see Supplementary Table 1).

Bidirectional communication between the sleep room and the control room was maintained through a speaker, microphone, and infrared camera, which allowed the experimenter to monitor and communicate with participants as needed. After participants woke up, resting-state EEG was again recorded in EO and EC conditions while participants lay on the bed. Participants then conducted again the PAL task.

### Statistical analyses Physiological data analysis

The data were analyzed via R and Python programming languages. Statistical analysis was conducted on EEG data from predetermined frontotemporal channels (F5, F6, FC3, FC4, FC5, FC6, C5, C6, C3, C4, F7, F8, FT7, FT8, T7, T8, TP7, TP8, CP5, CP6, CP3, CP4) across both the target stimulation frequency (0.75 Hz) and the entire delta frequency band (0.5-3.9 Hz). To assess the effects of the stimulation protocol on oscillatory activity, baseline normalization was performed by dividing post-stimulation power during resting-state by pre-stimulation power in the EC condition for each electrode. This normalization was conducted separately for each resting-state measurement timepoint (before and after sleep; see Figure 1) and each sleep stage (N1, N2, N3 and REM [Rapid Eye Movement] sleep). Baseline delta power (0.5-3.9 Hz) was analyzed to confirm that there were no significant differences between stimulation conditions. No significant differences were found, indicating that the baseline delta power can be validly used as a reference to assess changes in delta power during sleep and resting states following stimulation (see Supplementary Table S2).

The baseline-normalized data were analyzed using linear mixed-effects models, fitted with the *lmer* function from the *lme4* R package ^48^. Fixed effects included the stimulation protocol (*Protocol*, 2 levels), EEG measurement timepoint (*Timepoint*, 3 levels (baseline, resting state after intervention before sleep, after sleep)) for resting-state EEG, and sleep stage (*SleepStage*, 5 levels), EEG channel (*Electrode*, 22 levels) and their interactions. A by-participant random intercept term was included to account for the non-independence of observations within subjects. The significance of the fixed effects in the model was assessed using Satterthwaite’s approximation for degrees of freedom and F-tests from the *anova* function in the *lmer Test* R package ^49^. The critical significance level (alpha error) for all tests was set at 0.05. When significant effects were found, post-hoc pairwise comparisons were conducted using the estimated marginal means obtained from the model via the *emmeans* R package ^50^. These estimated means account for both random effects and the main factors in the model. The false discovery rate (FDR) was used to adjust for multiple comparisons. The decision to use estimated marginal means rather than pairwise t-tests for post-hoc testing was based on their ability to better reflect the structure of the linear mixed model, as they account for random effects and the dependencies within observations across multiple levels of factors. In addition, we conducted whole-brain analyses of the EEG at the sensor level during sleep.

These analyses contrasted delta frequency activity between stimulation and sham conditions for each sleep stage using permutation paired t-tests. The analysis involved 5000 permutations, and correction for multiple comparisons was applied using the FDR method. Furthermore, a statistical analysis was performed on global efficiency across all channels.

Similar to the power analysis, post-stimulation global efficiency during sleep was baseline-corrected by dividing it by pre-stimulation global efficiency in the EC condition for each sleep stage and participant. The baseline-normalized data were entered into a linear mixed model with the fixed effects sitmulation protocol (*Protocol,* 2 levels), sleep stage (*SleepStage*, 5 levels) and their interactions. Post-hoc pairwise comparisons were conducted using the estimated marginal means obtained from the linear mixed-effects model, with FDR correction applied for multiple comparisons.

For spindle activity, the mean fast and slow spindle counts were calculated across frontal and parietal channels for each participant, separately for N2 and N3 sleep stages and stimulation conditions. Paired t-tests were conducted to compare mean spindle counts between stimulation conditions in N2 and N3. P-values were adjusted using the FDR correction.

For sleep efficiency, the ratio of each sleep stage, and sleep onset latency, paired t-tests were performed to assess differences between stimulation conditions. Sleep efficiency was calculated as the total sleep time divided by the total time in bed, expressed as a percentage. The ratio of each sleep stage (N1, N2, N3 and REM) was calculated by dividing the time spent in each stage by the total sleep time for each participant. Sleep onset latency was defined as the time measured from lights off to the onset of the first sleep stage. Paired t-tests were conducted separately to compare these parameters between stimulation conditions, with P-values adjusted for multiple comparisons using the FDR correction.

### Behavioral data analysis

To compare PAL task performance between the stimulation and sham conditions, test scores were analyzed using a two-way repeated measures ANOVA, conducted with the *ezANOVA* function in R, and the results are summarized in Table 6. The ANOVA model included two factors: Condition (two levels: stimulation and sham), Time (two levels: pre-and post-sleep) and their interaction.

## Results

### Tolerability of stimulation protocols

Overall, no severe side effects were observed, and side effects reported during and after stimulation were only transient. Descriptive statistics of the presence and intensity of side effects can be found in the supplementary materials (Supplementary Table S3).

We conducted one-way repeated-measures ANOVAs to assess potential differences of side effects between the intervention groups. A significant effect was found for reports of tingling (F (1, 15) = 6.96, p = 0.02) and pain sensations (F (1, 15) = 9.52, p = 0.01) during stimulation, with larger tingling and pain scores corresponding to the trough-synchronized rTMS+tACS protocol (Table 1 and Supplementary Table S3). To assess possible effects of tingling and pain on delta frequency activity in the trough-synchronized rTMS+tACS condition, we performed Spearman’s rank correlation analyses. Specifically, the correlation between delta frequency activity (0.75 and 0.5-3.9 Hz) recorded after stimulation (Before Sleep) and tingling/pain scores was tested. The analyses showed no significant correlation between the intensity of tingling/pain and delta frequency activity, suggesting that these sensations did not affect delta activity (Supplementary Table S4). No other side effects showed significant differences across stimulation protocols, as outlined in Table 1.

**Table 1.**
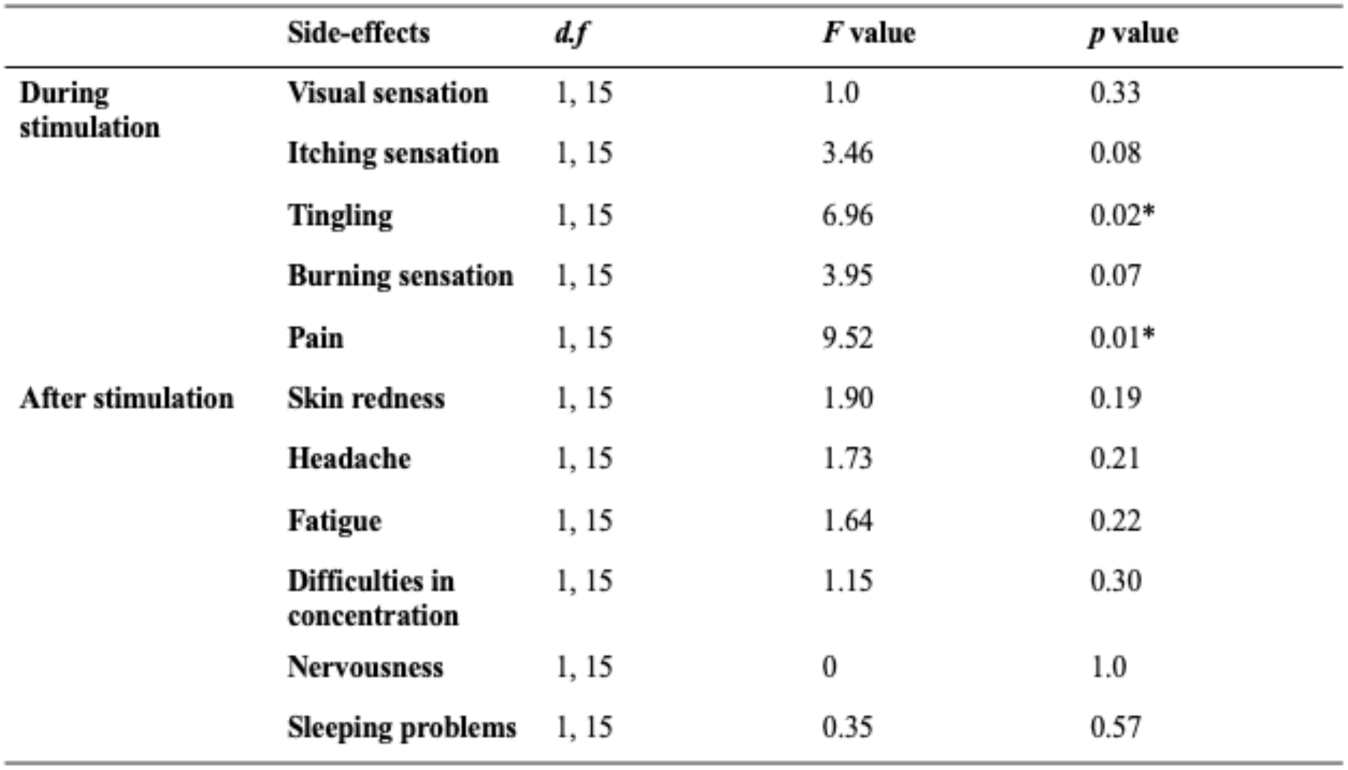
The presence and intensity of stimulation-related side effects was analyzed by one-way repeated-measures ANOVAs. Detailed ratings of the presence and intensity of side effects are documented in Table S3 of the supplementary materials. d.f. = degrees of freedom. Asterisks indicate significant differences.

|  | Side-effects | *df* | *F* value | *p* value |
| --- | --- | --- | --- | --- |
| **During stimulation** | **Visual sensation** | 1, 15 | 1.0 | 0.33 |
|  | **Itching sensation** | 1, 15 | 3.46 | 0.08 |
|  | **Tingling** | 1, 15 | 6.96 | 0.02* |
|  | **Burning sensation** | 1, 15 | 3.95 | 0.07 |
|  | **Pain** | 1, 15 | 9.52 | 0.01* |
| **After stimulation** | **Skin redness** | 1, 15 | 1.90 | 0.19 |
|  | **Headache** | 1, 15 | 1.73 | 0.21 |
|  | **Fatigue** | 1, 15 | 1.64 | 0.22 |
|  | **Difficulties in concentration** | 1, 15 | 1.15 | 0.30 |
|  | **Nervousness** | 1, 15 | 0 | 1.0 |
|  | **Sleeping problems** | 1, 15 | 0.35 | 0.57 |

### Spectrum analysis of resting-state EEG in the delta frequency range

The linear mixed model analyses conducted to assess stimulation-induced alterations of brain oscillations at 0.75 Hz revealed significant main effects of Protocol (EO: F(1, 1637.3) =24.67, p < 0.001, EC: F(1, 1678.9) = 6.15, p < 0.001), Timepoint (EO: F(2, 1637.8) = 61.20, p < 0.001, EC: F(2, 1671.9) = 26.84, p < 0.001), Channel (EO: F(21, 1632.1) = 2.15, p = 0.002, EC: F(21, 1669.8) = 1.82, p = 0.01) and the Protocol * Timepoint interactions in the EO condition (F(2, 1633.0) = 27.59, p < 0.001), as summarized in Table 2 and Figure 3.

**Figure 3.**
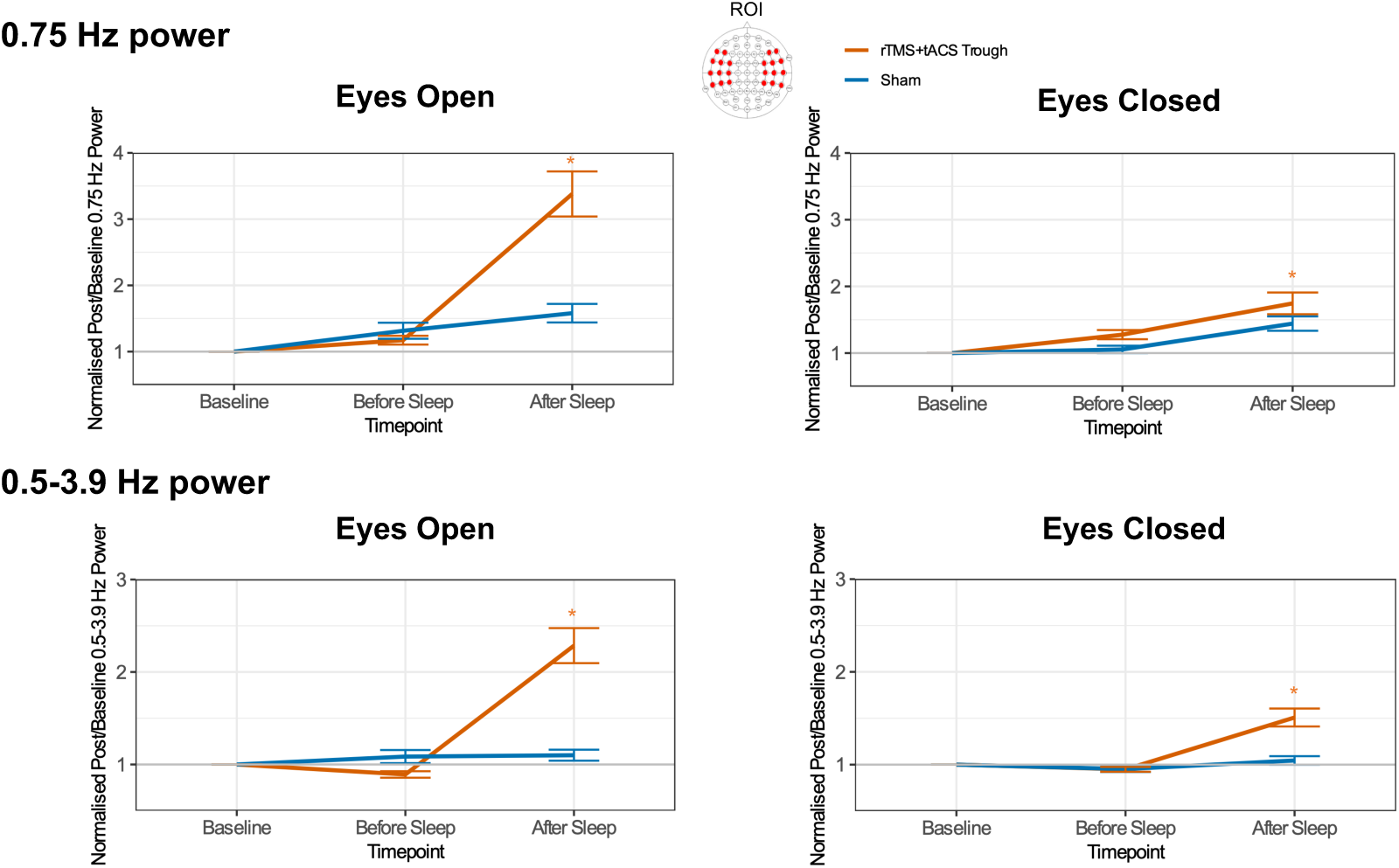
Changes in baseline-normalized power of the frontotemporal 0.75 Hz and the entire delta frequency band (0.75 Hz and 0.5-3.9 Hz) activity as a function of stimulation protocol and timepoint in resting-state EEG. Asterisks indicate significant pair-wise differences relative to sham (p < 0.05). Error bars represent the standard error of means.

**Table 2.**
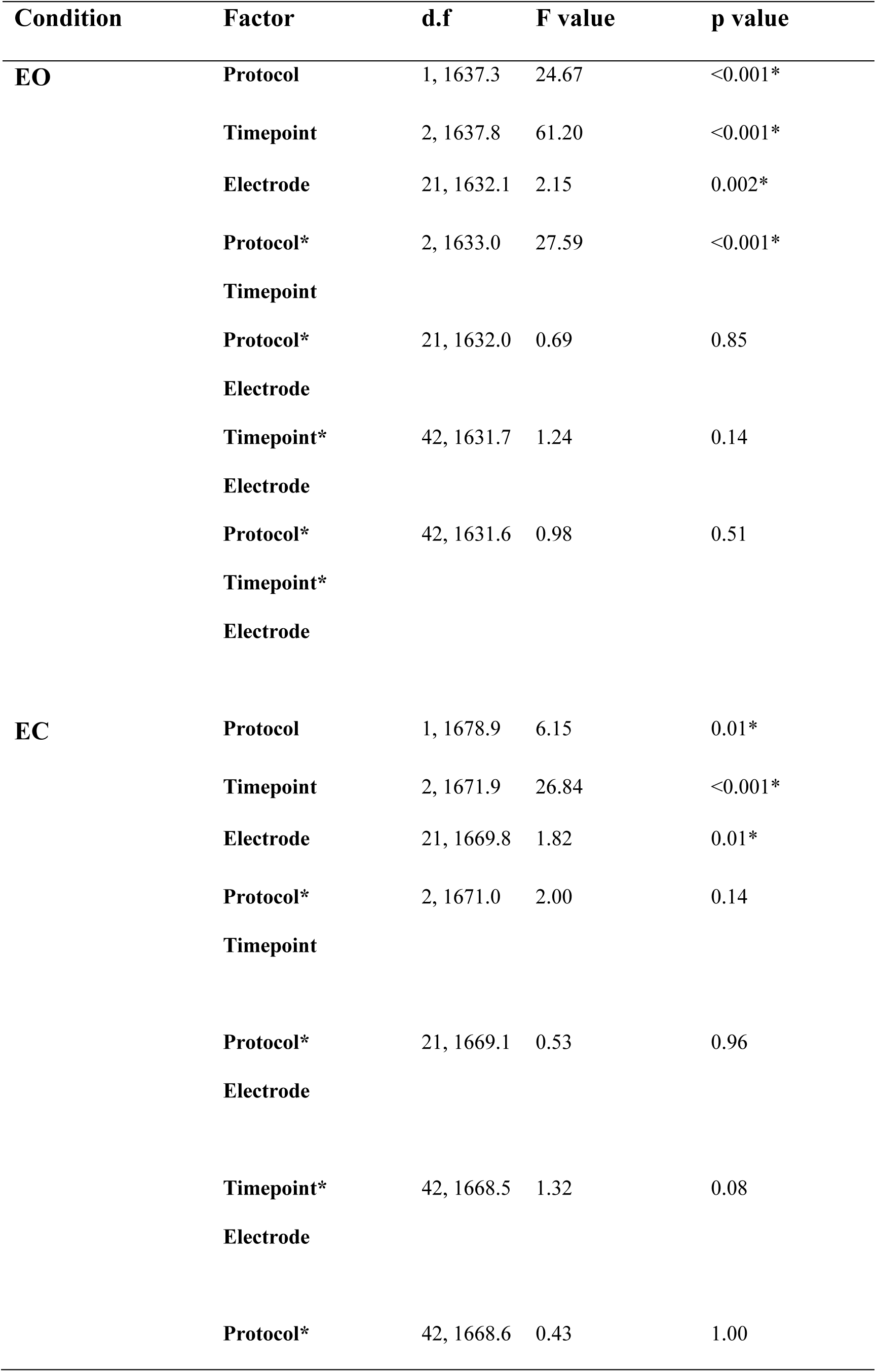
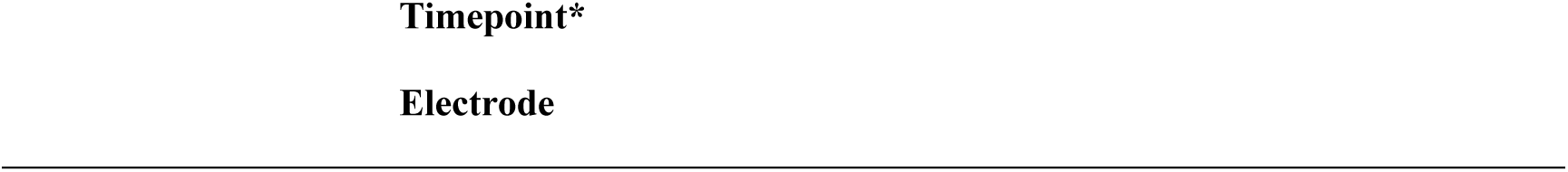
Results of the effect analysis performed by linear mixed models conducted for baseline-normalized 0.75 Hz activity during the resting state. EO indicates Eyes Open and EC indicates Eyes Closed conditions. d.f. = degrees of freedom. Asterisks indicate significant differences.

Subsequent post-hoc comparisons revealed significantly higher 0.75 Hz activity following sleep in the real stimulation condition compared to sham in both, EO and EC conditions (Supplementary Table S5).

We furthermore performed linear mixed model analyses on normalized oscillations of the whole delta frequency band (0.5-3.9 Hz). These showed significant main effects of Protocol (EO: F(1, 1631.1) = 31.38, p < 0.001, EC: F(1, 1733.9) = 18.04, p < 0.001), Timepoint (EO: F(2, 1632.6) = 53.19, p < 0.001, EC: F(2, 1731.1) = 31.52, p < 0.001), and a significant Protocol * Timepoint interaction (EO: F(2, 1627.8) = 46.95, p < 0.001, EC: F(2, 1731.2) = 18.43, p < 0.001). Also these effects were observed in both, EO and EC conditions, as shown in Table 3 and Figure 3.

**Table 3.**
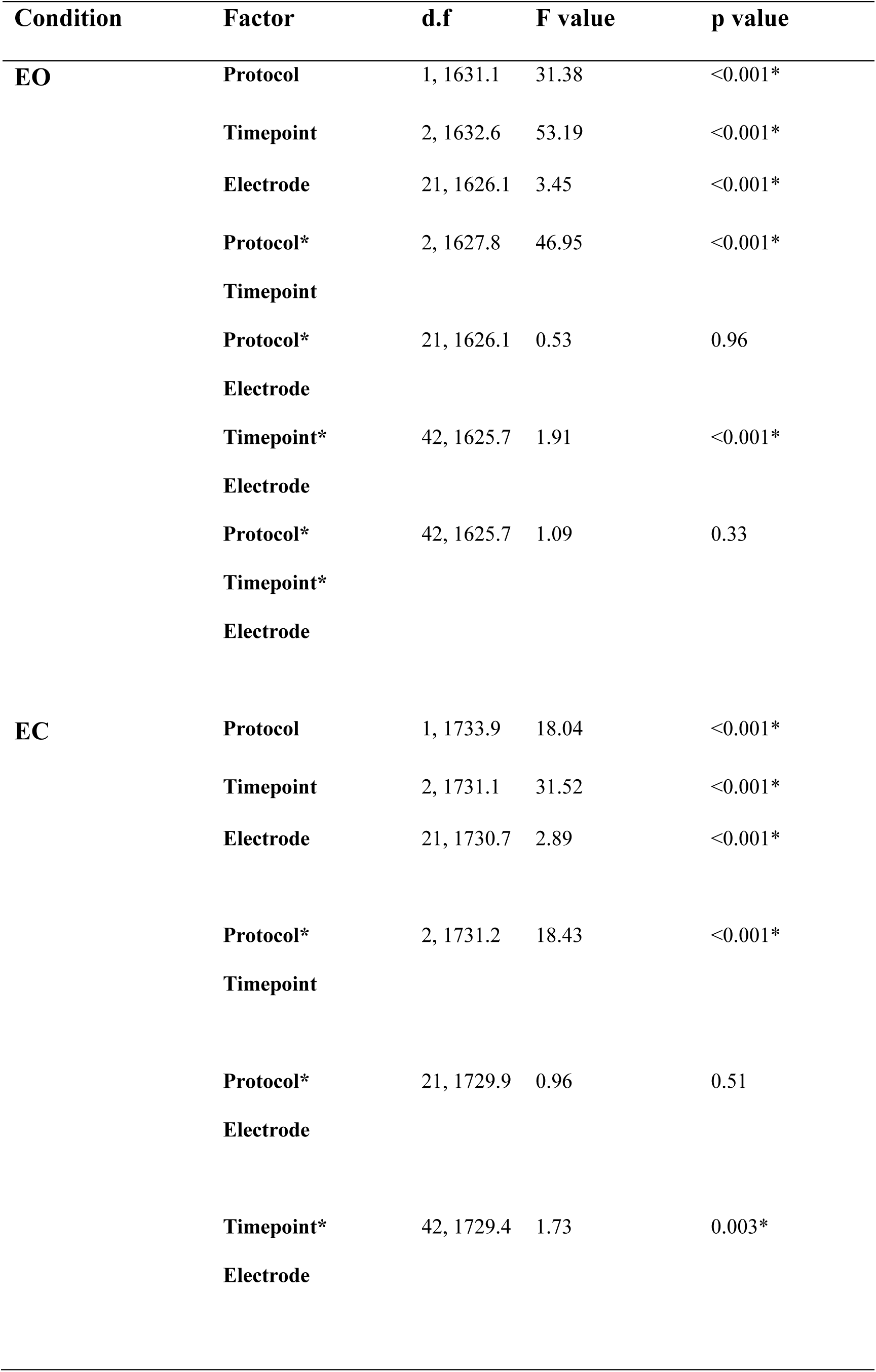
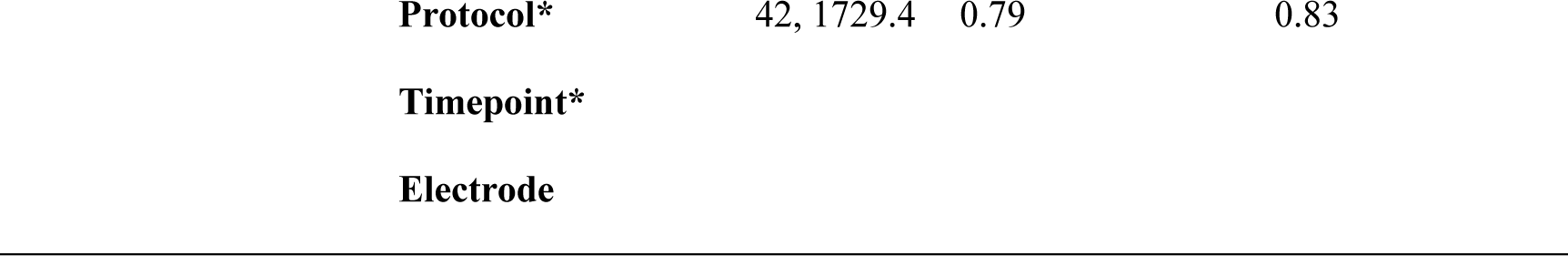
Results of the effect analysis performed by linear mixed models for baseline-normalized entire delta frequency activity (0.5-3.9 Hz) during the resting state. EO indicates Eyes Open and EC indicates Eyes Closed conditions. d.f. = degrees of freedom. Asterisks indicate significant differences.

| Condition | Factor | d.f | F value | p value |
| --- | --- | --- | --- | --- |
| **EO** | **Protocol** | 1, 1631.1 | 31.38 | <0.001* |
|  | **Timepoint** | 2, 1632.6 | 53.19 | <0.001* |
|  | **Electrode** | 21, 1626.1 | 3.45 | <0.001* |
|  | **Protocol\*** | 2, 1627.8 | 46.95 | <0.001* |
|  | **Timepoint** |  |  |  |
|  | **Protocol\*** | 21, 1626.1 | 0.53 | 0.96 |
|  | **Electrode** |  |  |  |
|  | **Timepoint\*** | 42, 1625.7 | 1.91 | <0.001* |
|  | **Electrode** |  |  |  |
|  | **Protocol\*** | 42, 1625.7 | 1.09 | 0.33 |
|  | **Timepoint\*** |  |  |  |
|  | **Electrode** |  |  |  |
| **EC** | **Protocol** | 1, 1733.9 | 18.04 | <0.001* |
|  | **Timepoint** | 2, 1731.1 | 31.52 | <0.001* |
|  | **Electrode** | 21, 1730.7 | 2.89 | <0.001* |
|  | **Protocol\*** | 2, 1731.2 | 18.43 | <0.001* |
|  | **Timepoint** |  |  |  |
|  | **Protocol\*** | 21, 1729.9 | 0.96 | 0.51 |
|  | **Electrode** |  |  |  |
|  | **Timepoint\*** | 42, 1729.4 | 1.73 | 0.003* |
|  | **Electrode** |  |  |  |

|  |  |  |  |
| --- | --- | --- | --- |
| **Protocol\*** | 42, 1729.4 | 0.79 | 0.83 |
| **Timepoint\*** |  |  |  |
| **Electrode** |  |  |  |

Post-hoc comparisons revealed significantly larger 0.5-3.9 Hz activity following sleep in the real stimulation condition compared to sham, in both EO and EC conditions (Supplementary Table S6).

### Spectral changes in delta oscillatory activity across sleep stages

The linear mixed model analyses conducted to assess stimulation-induced alternations of brain oscillations at 0.75 and 0.5-3.9 Hz showed significant main effects of Sleep Stage (0.75 Hz: F(4, 2845.5) = 886.4, p < 0.001, 0.5-3.9 Hz: F(4, 2946.4) = 18.04, p < 0.001), Channel (0.75 Hz: F(21, 2844.6) = 2.96, p < 0.001, 0.5-3.9 Hz: F(21, 2945.3) = 10.51, p < 0.001), and a significant Protocol * Sleep Stage interaction (0.75 Hz: F(4, 2845.2) = 4.03, p = 0.003, 0.5-3.9 Hz: F(4, 2946.1) = 9.97, p < 0.001), as shown in Table 4 and Figure 4. Subsequent post-hoc comparisons revealed significantly larger 0.75 and 0.5-3.9 Hz activity in the N3 sleep stage in the real stimulation condition compared to sham (Supplementary Table S7 and S8).

**Figure 4.**
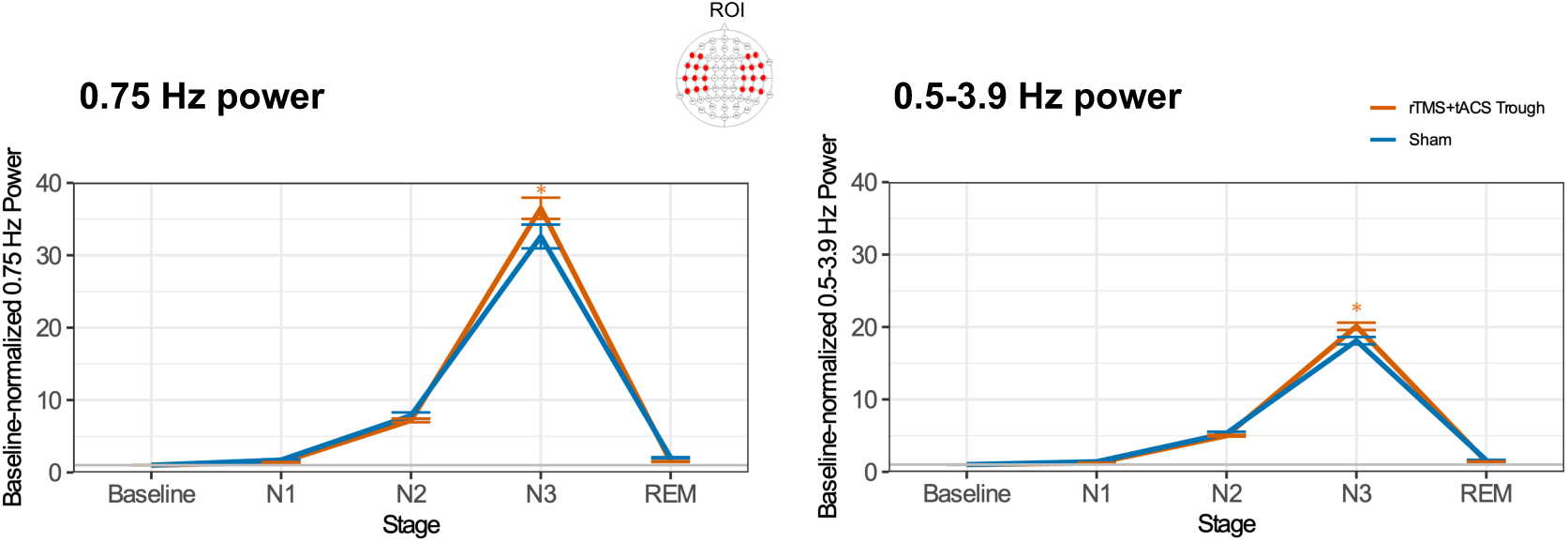
Changes in baseline-normalized power of the frontotemporal 0.75 Hz and the entire delta frequency band (0.75 Hz and 0.5-3.9 Hz) activity as a function of stimulation protocol and sleep stage. Asterisks indicate significant pair-wise differences relative to sham (p < 0.05). Error bars represent the standard error of means.

**Table 4.**
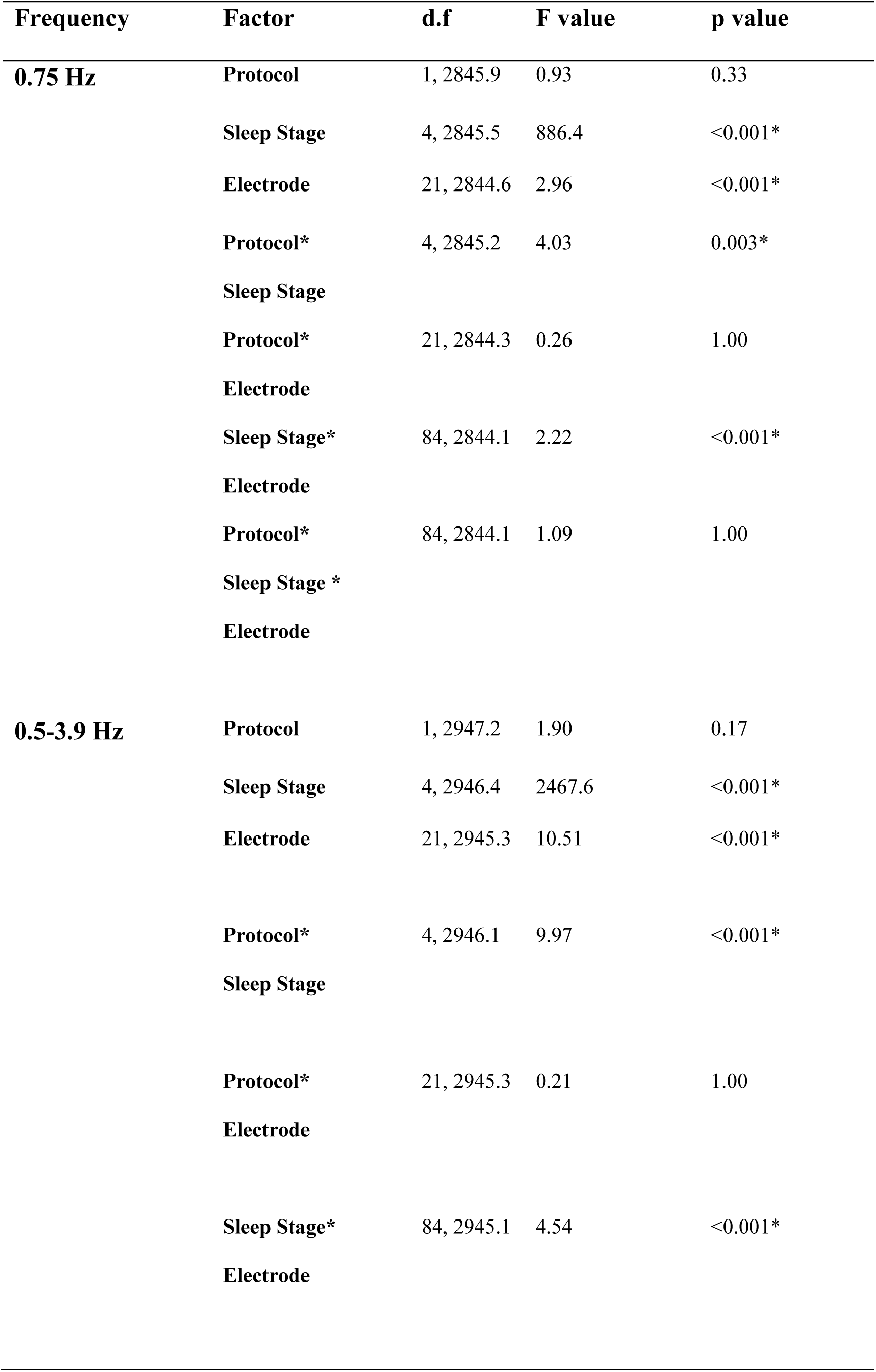
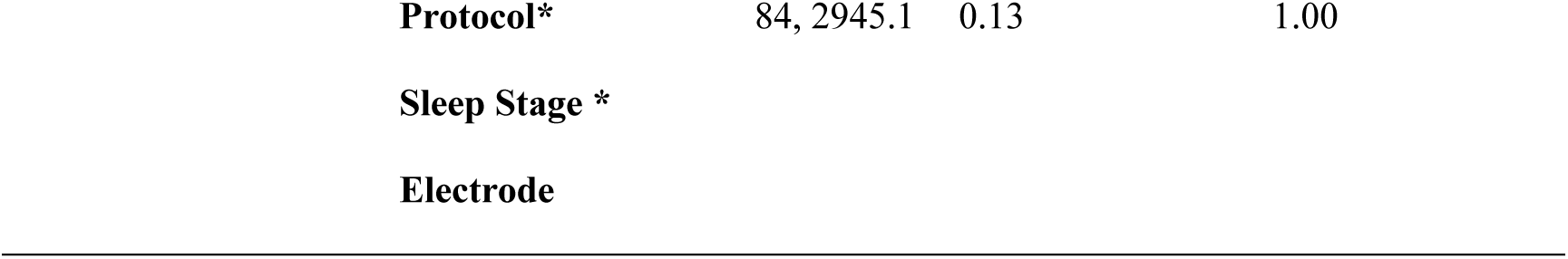
Results of the effect analysis performed by linear mixed models for baseline-normalized 0.75 Hz activity and entire delta frequency activity (0.5-3.9 Hz) during sleep. d.f. = degrees of freedom. Asterisks indicate significant differences.

| Frequency | Factor | d.f | F value | p value |
| --- | --- | --- | --- | --- |
| **0.75 Hz** | **Protocol** | 1, 2845.9 | 0.93 | 0.33 |
|  | **Sleep Stage** | 4, 2845.5 | 886.4 | <0.001* |
|  | **Electrode** | 21, 2844.6 | 2.96 | <0.001* |
|  | **Protocol\*** | 4, 2845.2 | 4.03 | 0.003* |
|  | **Sleep Stage** |  |  |  |
|  | **Protocol\*** | 21, 2844.3 | 0.26 | 1.00 |
|  | **Electrode** |  |  |  |
|  | **Sleep Stage\*** | 84, 2844.1 | 2.22 | <0.001* |
|  | **Electrode** |  |  |  |
|  | **Protocol\*** | 84, 2844.1 | 1.09 | 1.00 |
|  | **Sleep Stage \*** |  |  |  |
|  | **Electrode** |  |  |  |
| **0.5-3.9 Hz** | **Protocol** | 1, 2947.2 | 1.90 | 0.17 |
|  | **Sleep Stage** | 4, 2946.4 | 2467.6 | <0.001* |
|  | **Electrode** | 21, 2945.3 | 10.51 | <0.001* |
|  | **Protocol\*** | 4, 2946.1 | 9.97 | <0.001* |
|  | **Sleep Stage** |  |  |  |
|  | **Protocol\*** | 21, 2945.3 | 0.21 | 1.00 |
|  | **Electrode** |  |  |  |
|  | **Sleep Stage\*** | 84, 2945.1 | 4.54 | <0.001* |
|  | **Electrode** |  |  |  |

|  |  |  |  |
| --- | --- | --- | --- |
| **Protocol\*** | 84, 2945.1 | 0.13 | 1.00 |
| **Sleep Stage \*** |  |  |  |
| **Electrode** |  |  |  |

While whole-brain, sensor-level analyses did not reveal significant channel-level changes after applying multiple comparison correction (FDR), a general trend of increased delta power across N2 and N3 stages was observed for both delta band categories, when contrasted against sham (p < 0.05, corrected), as shown in Figure 5.

**Figure 5.**
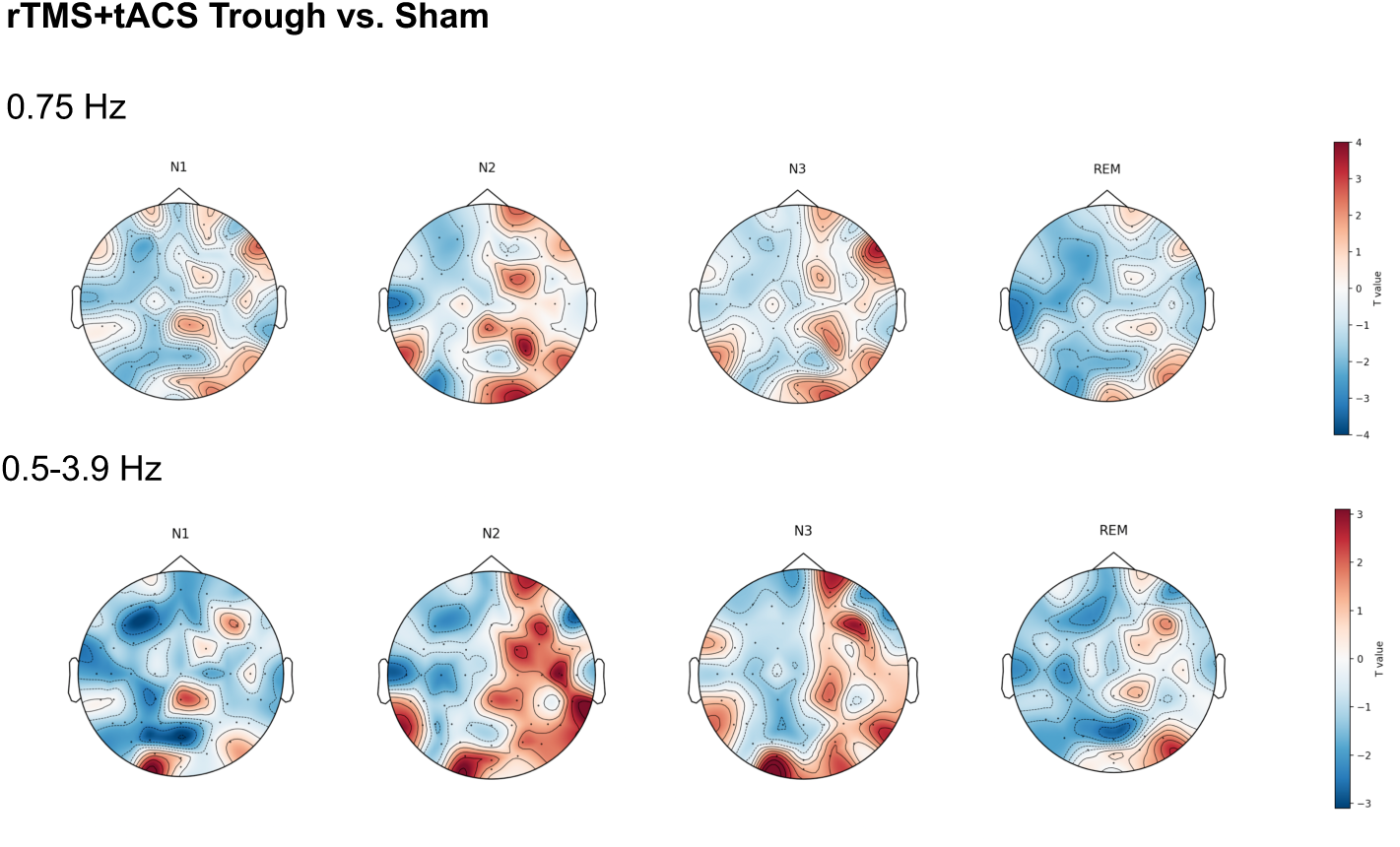
Whole-brain, sensor level t-contrasts comparing baseline-corrected changes in delta activity (0.75 Hz and 0.5-3.9 Hz) between real stimulation and sham stimulation are shown. The maps visualize the differential distribution of changes in delta power in paired comparisons, revealing a broader distribution of enhanced delta power in the N2 and N3 sleep stages compared to the N1 and REM sleep stages. This was consistently observed across both delta frequency ranges.

### Changes in global functional network efficiency of resting-state EEG

The linear mixed model analyses conducted to assess stimulation-induced changes in functional connectivity at 0.75 Hz revealed a significant main effect of Protocol in the EO condition (F(1, 69.8) = 6.78, p = 0.01) and a significant main effect of Timepoint in the EC condition (F(2, 67.1) = 3.57, p = 0.03), as summarized in Table 5 and Figure 6. Subsequent post-hoc comparisons revealed significantly lower global efficiency following sleep in the real stimulation condition compared to sham in the EO condition (Supplementary Table S9). Within the 0.5-3.9 Hz frequency range, we observed a significant main effect for Protocol (F(1, 67.5) = 9.80, p = 0.003) and a significant interaction of Protocol and Timepoint (F(2, 64.5) = 3.84, p = 0.03) in the EO condition, as summarized in Table 6 and Figure 6. Post-hoc comparisons revealed significantly lower global efficiency before sleep in the real stimulation condition compared to sham in the EO condition (Supplementary Table S10).

**Figure 6.**
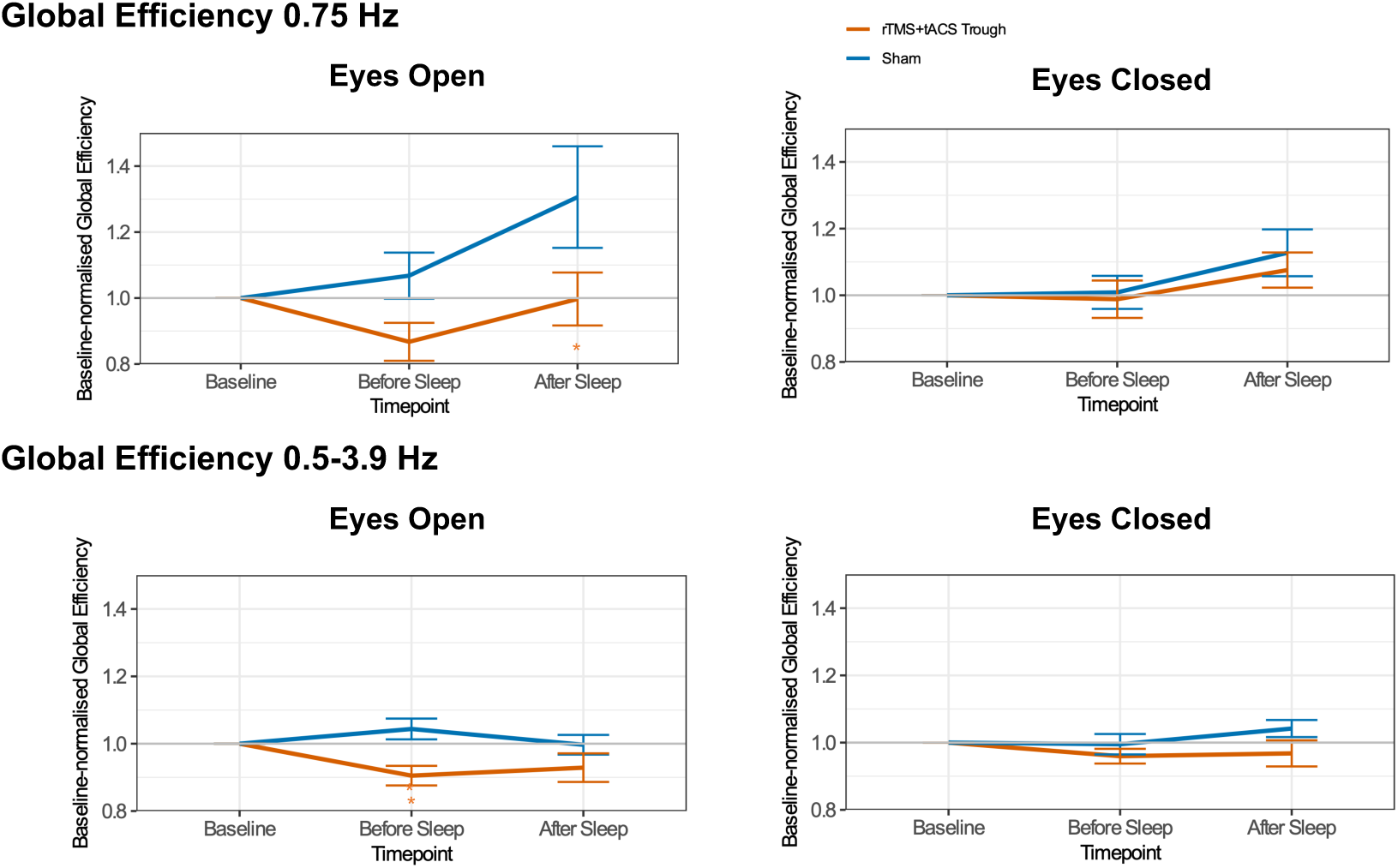
Changes in baseline-normalized global functional network efficacy in the delta frequency range (0.75 Hz and 0.5-3.9 Hz) as a function of stimulation protocol and timepoint in resting-state EEG. Asterisks indicate significant pair-wise differences relative to sham (p < 0.05). Error bars represent the standard error of means.

**Table 5.** Results of the effect analysis performed by linear mixed models for baseline-normalized global efficiency at 0.75 Hz during the resting state. d.f. = degrees of freedom. Asterisks indicate significant differences.

| **Condition** | **Factor** | **d.f** | **F value** | **p value** |
| --- | --- | --- | --- | --- |
| **EO** | **Protocol** | 1, 69.8 | 6.78 | 0.01* |
|  | **Timepoint** | 2, 69.6 | 3.05 | 0.05 |
|  | **Protocol\*** | 2, 70.1 | 2.10 | 0.13 |
|  | **Timepoint** |  |  |  |
| **EC** | **Protocol** | 1, 69.9 | 0.62 | 0.43 |
|  | **Timepoint** | 2, 67.1 | 3.57 | 0.03* |
|  | **Protocol\*** | 2, 67.3 | 0.21 | 0.82 |
|  | **Timepoint** |  |  |  |

### Changes in global functional network efficiency across sleep stages

The linear mixed model analyses revealed a significant main effect of Sleep Stage on global efficiency in both delta frequency ranges (0.75 Hz: F(4, 126.3) = 14.35, p < 0.001, 0.5-3.9 Hz: F(4, 127.8) = 74.43, p < 0.001). No main effect of condition or interaction between condition and sleep stage was detected (Table 7). Given the importance of discovering the sleep-stage-dependent effects of stimulation on functional connectivity, we conducted exploratory post hoc pairwise comparisons to investigate potential trends within each sleep stage for each condition. These comparisons revealed higher global efficiency in the real stimulation condition compared to sham, specifically within the N2 sleep stage in both delta frequency ranges (Supplementary Table S11, S12 and Figure 7).

**Figure 7.**
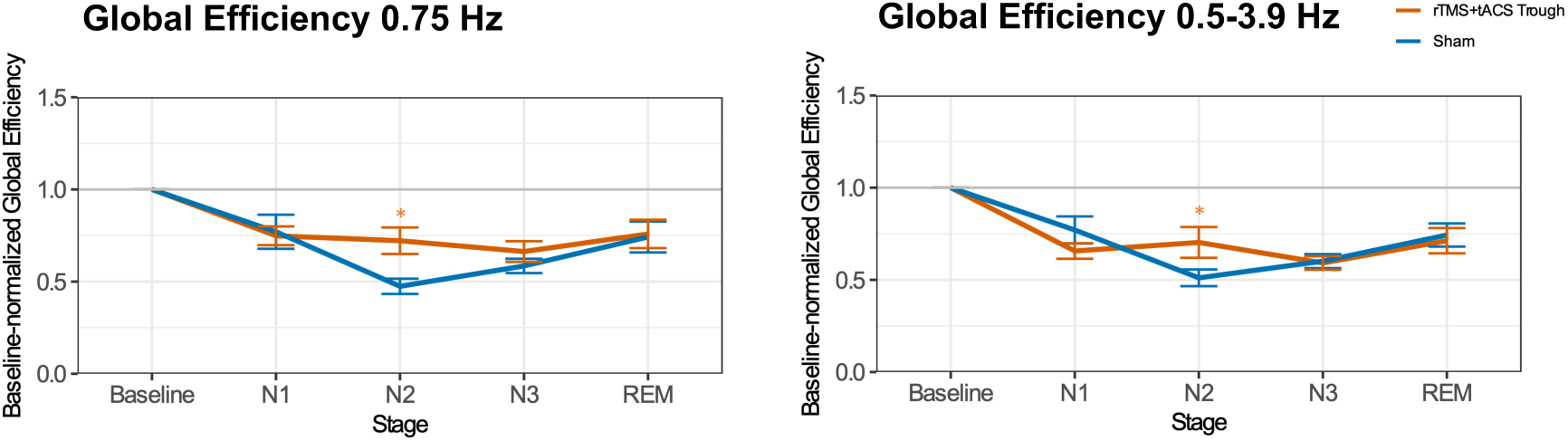
Changes in baseline-normalized global functional network efficiency in the delta frequency range (0.75 Hz and 0.5-3.9 Hz) as a function of stimulation protocol and sleep stage. Asterisks indicate significant pair-wise differences relative to sham (p < 0.05). Error bars represent the standard error of means.

**Table 6.** Results of the effect analysis performed by linear mixed models for baseline-normalized global efficiency across the entire delta frequency range (0.5-3.9 Hz) during the resting state. d.f. = degrees of freedom. Asterisks indicate significant differences.

| **Condition** | **Factor** | **d.f** | **F value** | **p value** |
| --- | --- | --- | --- | --- |
| **EO** | **Protocol** | 1, 67.5 | 9.80 | 0.003* |
|  | **Timepoint** | 2, 64.6 | 1.28 | 0.29 |
|  | **Protocol\*** | 2, 64.5 | 3.84 | 0.03* |
|  | **Timepoint** |  |  |  |
| **EC** | **Protocol** | 1, 66.8 | 3.54 | 0.06 |
|  | **Timepoint** | 2, 63.1 | 1.05 | 0.36 |
|  | **Protocol\*** | 2, 63.2 | 1.63 | 0.20 |
|  | **Timepoint** |  |  |  |

**Table 7.** Results of the effect analysis performed by linear mixed models for baseline-normalized global efficiency at 0.75 Hz and across the entire delta frequency range (0.5-3.9 Hz) during sleep. d.f. = degrees of freedom. Asterisks indicate significant differences.

| **Frequency** | **Factor** | **d.f** | **F value** | **p value** |
| --- | --- | --- | --- | --- |
| **0.75 Hz** | **Protocol** | 1, 130.5 | 2.74 | 0.10 |
|  | **Sleep Stage** | 4, 126.3 | 14.35 | <0.001* |
|  | **Protocol\*** | 4, 126.3 | 1.59 | 0.18 |
|  | **Sleep Stage** |  |  |  |
| **0.5-3.9 Hz** | **Protocol** | 1, 127.9 | 0.004 | 0.95 |
|  | **Sleep Stage** | 4, 127.8 | 74.43 | <0.001* |
|  | **Protocol\*** | 4, 128.1 | 2.43 | 0.051 |
|  | **Sleep Stage** |  |  |  |

### Spindle counts

Paired t-tests revealed no significant differences in fast or slow spindle counts between stimulation conditions in either frontal or parietal channels across N2 and N3 sleep stages (Figure 8 and Supplementary Table S13).

**Figure 8.**
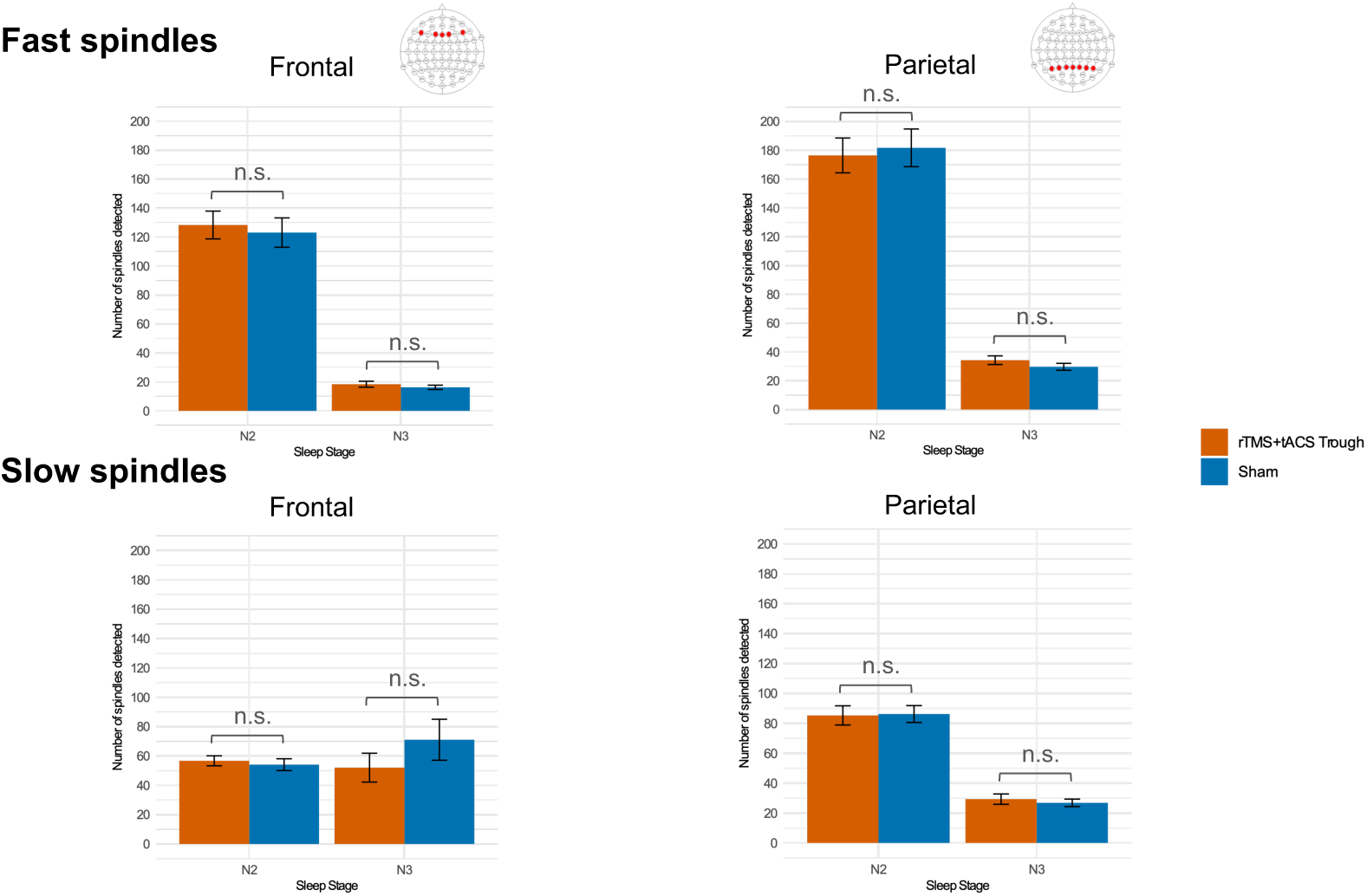
The mean counts of fast (12-15 Hz) and slow (9-11.9 Hz) spindles in frontal and parietal channels during N2 and N3 sleep stages in real stimulation and sham conditions. Error bars represent the standard error of means.

### Sleep stage ratios, sleep onset latency and sleep efficiency

Paired t-tests revealed no significant differences of each sleep stage ratios, sleep onset latency and sleep efficiency between the stimulation conditions (Figure 9 and Supplementary Table S14, 15).

**Figure 9.**
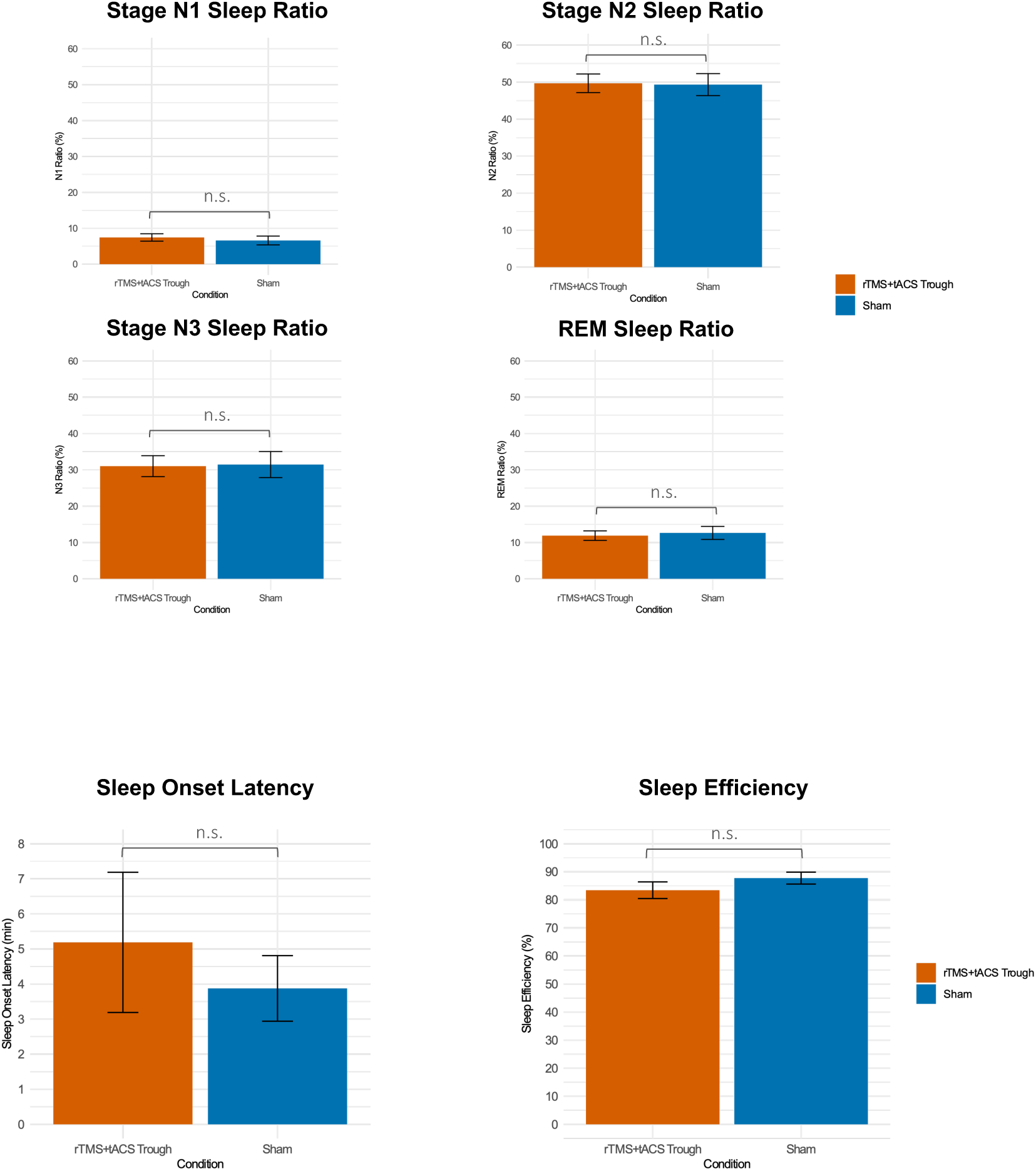
The mean ratios of each sleep stage, sleep onset latency, and sleep efficiency. Error bars represent the standard error of means.

### PAL task

The results of the repeated measures ANOVA revealed only a significant main effect of Time (F(1, 15)=13.86, p=0.002), but no significant effect of Condition (F(1, 15)=0.03, p=0.87), indicating that there was an overall improvement in memory performance after sleep in both conditions. However, memory performance did not differ significantly between the stimulation and sham conditionns, and no interaction between the conditions and time points was found (Table 8). Figure 10 shows the task performance for each condition.

**Figure 10.**
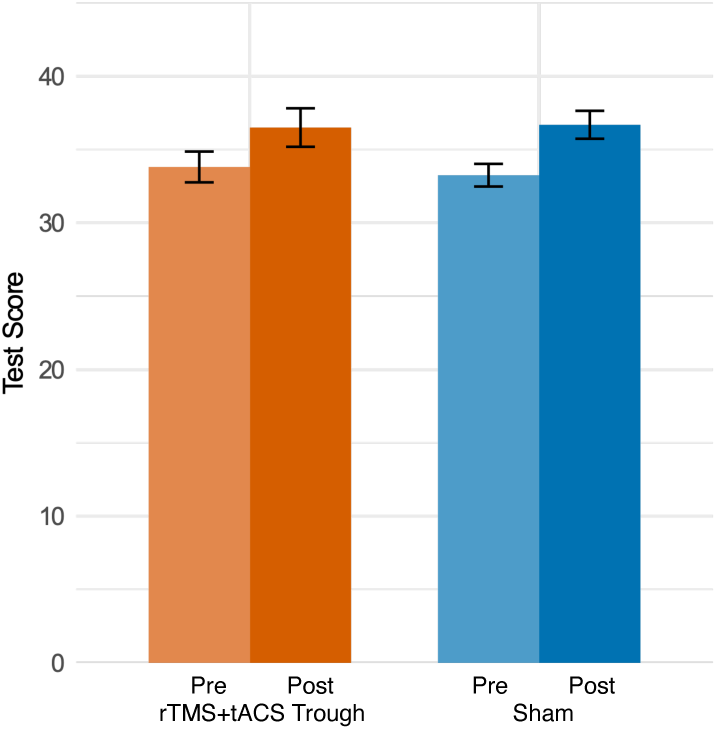
PAL task scores before and after stimulation in the real and sham conditions. Error bars represent the standard error of means.

**Table 8.** Results of the effect analysis performed by a two-way repeated measures ANOVA on the test scores for the paired associate learning task. d.f. = degrees of freedom. Asterisks indicate significant differences.

| **Factor** | **d.f.** | **F value** | **P value** |
| --- | --- | --- | --- |
| **Condition** | 1, 15 | 0.03 | 0.87 |
| **Timepoint** | 1, 15 | 13.86 | 0.002* |
| **Condition\*** | 1, 15 | 0.45 | 0.51 |
| **Timepoint** |  |  |  |

## Discussion

The polysomnographic findings demonstrated that trough-synchronized rTMS and tACS, previously shown to induce and stabilize delta oscillatory activity during wakefulness ^28^, induced sleep stage-specific effects. Specifically, trough-synchronized rTMS and tACS led to a significant increase of delta activity at the stimulation frequency of 0.75 Hz and across the entire delta frequency range (0.5-3.9 Hz) during the N3 sleep stage compared to the sham condition, whereas no significant changes in delta activity were observed during N1, N2, or REM sleep stages. A sustained enhancement in delta activity was observed in resting-state EEG even after sleep in the stimulation condition. In terms of functional connectivity during sleep, measured by global efficiency, stimulation led to higher global efficiency during the N2 sleep stage compared to sham.

In contrast to previous findings, where stimulation led to an immediate increase in delta oscillatory activity ^28^, the current study did not show such an enhancement before sleep. One key difference between these studies is that participants in the present study underwent sleep deprivation prior to stimulation. While sleep deprivation is known to increase delta EEG activity, particularly during subsequent sleep ^51–53^, its effects on EEG activity during wakefulness are not well understood. Moreover, functional connectivity in the eyes-open resting state decreased following stimulation, which contrasts with previous findings of increased connectivity. It is possible that sleep deprivation affected the responsiveness of delta frequency oscillatory activity during subsequent wakefulness. Further reseach is warranted to clarify how sleep deprivation interacts with brain stimulation protocols.

Regarding sleep statistics, no significant differences were found in sleep stage ratios (N1, N2, N3 and REM), sleep onset latency, or overall sleep efficiency between stimulation and sham conditions. These findings indicate that the observed increase in delta activity was not attributable to differences in sleep stage proportions between conditions. Instead, the effects of stimulation specifically enhanced neuronal activity within the delta frequency range without altering sleep structure.

While the physiological results support our hypothesis that the combined rTMS and tACS protocol enhances delta activity and functional connectivity during, and delta activity also after sleep, our findings did not reveal a significant effect of rTMS and tACS on declarative memory performance. Memory performance of the PAL task improved overall after sleep in both conditions, but no significant difference was observed between real stimulation and sham conditions. These results differ from those of the previous study by Marshall et al. (2006), which showed that slow oscillating tDCS increased slow oscillations in the EEG and improved memory performance, accompanied by increased spindle activity, postulating a causal role of slow oscillations, sleep spindles, or both, on memory consolidation ^23^.

Our present findings that showed that stimulation enhanced delta oscillatory activity but left memory performance and spindle activity unchanged suggest that the enhancement of slow oscillatory activity alone is not sufficient to improve memory performance. This aligns with the concept that memory consolidation during sleep relies on the interplay between slow oscillations and sleep spindles, which facilitates hippocampal-neocortical transfer of memory ^54,55^. However, since the so-tDCS protocols introduced by Marshall et al. include also an LTP-like plasticity-inducing tDCS component, it cannto be ruled out that this protocol characteristic improved cognitive performance in the previous study.

A potential mechanistic explanation is that stimulation-induced slow oscillations did not mimic endogeneous slow oscillations relevant to memory consolidation during sleep. Prior studies have highlighted the variability of slow oscillatory activity during sleep, suggesting that different forms of slow oscillations may serve distinct functions. For instance, these distinguish between widespread, high-amplitude slow-waves and localized, lower-amplitude slow waves, the latter being linked to sleep spindles ^56,57^. Endogeneous slow oscillations alternate between up-states, characterized by increased neuronal activity and depolarization, and down-states, characterized by neuronal silence due to hyperpolarization ^58^. The up-state serves as a window for hippocampal-neocortical interactions ^59^ and triggers thalamocortical spindles ^60^. The coupling of slow oscillations and spindles has been proposed to facilitate synaptic plasticity ^61^, and spindles phase-locked to the up-state of slow oscillations have been associated with enhanced declarative memory performance ^62^. However, stimulation-induced slow oscillations may differ in their topographical characteristics or amplitude from endogeneous oscillations relevant for memory consolidation. These differences could alter their function, limiting their ability to trigger spindles and facilitate coupling, and as a result, hippocampal-neocortical transfer of memories may not occur.

Moreover, so-tDCS, as used in previous studies ^22,23^, induces also LTP-like plasticity ^24^, a mechanism not expected from the rTMS and tACS protocol employed in the current study. This difference in plasticity induction might explain why enhanced memory consolidation was observed following so-tDCS in previous studies but was absent in the current study.

Although stimulation in the current study did not enhance memory consolidation, this does not exclude other potential functions that the induced slow oscillations might serve. Beyond memory consolidation, slow oscillations during sleep are known to serve broader functions. For instance, disrupted slow-wave activity during NREM sleep has been associated with increased daytime sleep propensity, commonly observed in individuals with insomnia, which negatively impacts occupational functioning ^63–65^. Future studies should therefore examine whether stimulation-induced slow oscillations can restore slow-wave activity and mitigate symptoms associated with disrupted slow-wave activity in clinical populations.

Some limitations of the present study should be acknowledged. It cannot be excluded that the absence of enhanced memory performance may have resulted from the limited number of sleep cycles. Memory performance was assessed after three hours of sleep, whereas participants in the previous study had a full night sleep (11 pm until 6.30 am) ^23^. Studies have demonstrated a positive correlation between the number of NREM-REM sleep cycles and declarative memory performance ^66,67^. The reproducibility of the current findings under conditions of a full night sleep should be assessed to better capture the impact of the stimulation protocol on memory consolidation in future studies.

In conclusion, we demonstrated that rTMS synchronized to the trough of the tACS enhanced delta oscillatory activity in a sleep stage-specific manner. The present findings suggest that phase-synchronized rTMS and tACS, applied during wakefulness, can induce long-lasting modulation of specific oscillatory activity during sleep in humans. Given its sustained efficacy, this stimulation protocol holds potential for flexible application in future experimental designs. Furthermore, these results highlight the potential for clinical applications to address pathological slow-wave oscillations during sleep.

