## Supplementary Data for "Sleep stage-specific effects of 0.75 Hz phase-synchronized rTMS and tACS on delta frequency activity during sleep"

**Supplementary Information**

**Supplementary Table S1:**

For the first five participants, the study design involved a 3-hour time in bed, with participants being woken up 3 hours after the lights were turned off (lights-off group). For the remaining participants, total sleep time was maintained at 3 hours by waking them 3 hours after they entered the N1 sleep stage (N1 stage group). An independent t-test was performed to assess whether total sleep time differed between participants who were woken up three hours after lights-off and those who were woken up 3 hours after entering the N1 sleep stage. The analysis revealed no significant difference in total sleep time between the two groups, allowing for a combined analysis of all participants (p > 0.05). The results including the mean and standard errors (mean ± SE in minutes) are shown in the table below.

*Difference between groups*

|  | ***Mean ± SE***  ***(Lights off Group)*** | ***Mean ± SE***  ***(N1 Stage Group)*** | ***t value*** | ***p value*** |
| --- | --- | --- | --- | --- |
| **Total Sleep**  **Time** | 155.10 ± 5.38 | 156.89 ± 4.08 | -0.25 | 0.80 |

**Supplementary Table S2:**

To assess whether delta power (0.5-3.9 Hz) of resting-state EEG at baseline differed significantly between real and sham stimulation conditions, linear mixed model analyses were conducted with the fixed effect of stimulation protocol (2 levels) separately for the eyes open (EO) and eyes closed (EC) conditions. The means and standard errors (SE) are expressed in decibels (dB).

**0.75 Hz**

| **Condition** | ***Mean ± SE (dB)***  ***(Stimulation group)*** | ***Mean ± SE (dB)***  ***(Sham group)*** | ***d.f.*** | ***F value*** | ***p value*** |
| --- | --- | --- | --- | --- | --- |
| **EO** | -82.24 ± 0.62 | -79.26 ± 0.92 | 1, 15 | 4.45 | 0.05 |
| **EC** | -81.85 ± 0.60 | -81.14 ± 0.59 | 1, 15 | 1.68 | 0.21 |

**0.5-3.9 Hz**

| **Condition** | ***Mean ± SE (dB)***  ***(Stimulation group)*** | ***Mean ± SE (dB)***  ***(Sham group)*** | ***d.f.*** | ***F value*** | ***p value*** |
| --- | --- | --- | --- | --- | --- |
| **EO** | -86.27 ± 0.56 | -84.58 ± 0.82 | 1, 15 | 2.38 | 0.14 |
| **EC** | -86.46 ± 0.41 | -85.45 ± 0.54 | 1, 15 | 3.79 | 0.07 |

**Supplementary Table S3:**

Participants provided ratings for the presence and severity of side effects, which included assessments of visual sensations, itching, tingling, burning sensations and pain during stimulation, as well as skin redness, headache, fatigue, difficulty concentrating, nervousness, and sleep problems after stimulation. These side effects were evaluated using a numerical scale ranging from zero (indicating no sensation) to five (representing extremely strong sensations). The data are presented as mean values with standard deviations (mean ± SD). We conducted one-way repeated-measures ANOVAs to assess side effect differences between the intervention groups, and the results are depicted in Table 1 in the main text.

|  |  | **rTMS+tACS**  **Trough** | **Sham** |
| --- | --- | --- | --- |
| **During Stimulation** | Visual sensation | 0.41 ± 0.87 | 0.18 ± 0.53 |
|  | Itching sensation | 0.59 ± 0.94 | 0.24 ± 0.56 |
|  | Tingling | 2.06 ± 1.48 | 1.00 ± 1.12 |
|  | Burning sensation | 0.59 ± 1.23 | 0.00 ± 0.00 |
|  | Pain | 1.59 ± 1.54 | 0.29 ± 0.69 |
| **After**  **Stimulation** | Skin redness | 0.47 ± 1.28 | 0.29 ± 1.21 |
|  | Headache | 1.24 ± 1.68 | 0.71 ± 1.16 |
|  | Fatigue | 1.29 ± 1.31 | 1.06 ± 1.39 |
|  | Difficulties in concentration | 0.94 ± 1.24 | 0.71 ± 1.16 |
|  | Nervousness | 0.24 ± 0.56 | 0.24 ± 0.75 |
|  | Sleeping problems | 0.29 ± 0.77 | 0.47 ± 1.01 |

**Supplementary Table S4:**

To investigate whether the perceived tingling and pain, which were reported to be stronger in the trough-aligned rTMS and tACS condition than in the sham condition, affected delta activity in offline EEG recordings, we conducted Spearman's rank correlation analyses. These analyses tested the association between resting-state EEG delta activity (0.75 Hz and 0.5-3.9 Hz) recorded after stimulation (before sleep) and the tingling/pain scores. The results did not reveal a significant correlation between tingling/pain intensity and delta activity.

**0.75 Hz**

| **Sensastion** | **Condition** | ***Spearman***  ***Correlation***  ***Coefficient*** | ***p value*** |
| --- | --- | --- | --- |
| **Tingling** | **EO** | -0.39 | 0.15 |
|  | **EC** | 0.30 | 0.26 |
| **Pain** | **EO** | 0.01 | 0.98 |
|  | **EC** | -0.02 | 0.94 |

**0.5-3.9 Hz**

| **Sensastion** | **Condition** | ***Spearman***  ***Correlation***  ***Coefficient*** | ***p value*** |
| --- | --- | --- | --- |
| **Tingling** | **EO** | -0.16 | 0.56 |
|  | **EC** | -0.07 | 0.79 |
| **Pain** | **EO** | 0.02 | 0.94 |
|  | **EC** | -0.28 | 0.30 |

**Supplementary Table S5-6:**

We employed linear mixed models to assess the main effects and interactions of stimulation protocol (2 levels), timepoint (3 levels), EEG channel (22 levels) as within-subject factors on baseline-normalized delta power (0.75 Hz and 0.5-3.9 Hz) during the resting state.

In case of significant effects of these primary analyses (Table 2 and 3), post-hoc pairwise comparisons using the estimated marginal means approach were performed between stimulation protocols.

The analyses were conducted separately for the eyes open (EO) and eyes closed (EC) resting state conditions. Asterisks indicate significant differences.

Supplementary Table S5: Results of post-hoc pairwise contrasts between rTMS and tACS Trough and sham stimulation protocols for baseline-normalized 0.75Hz activity during the resting state are presented below. The table also shows the estimated marginal means and standard errors (mean ± SE) of baseline-normalized 0.75 Hz power of each stimulation condition.

*Before Sleep*

| Condition | *Estimated Marginal Mean ± SE*  *rTMS+tACS Group* | *Estimated Marginal Mean ± SE*  *Sham Group* | *t value* | *p value* |
| --- | --- | --- | --- | --- |
| **EO** | 1.16 ± 0.27 | 1.22 ± 0.27 | -0.29 | 0.77 |
| **EC** | 1.26 ± 0.12 | 1.04 ± 0.12 | 1.87 | 0.06 |

*After Sleep*

| Condition | *Estimated Marginal Mean ± SE*  *rTMS+tACS Group* | *Estimated Marginal Mean ± SE*  *Sham Group* | *t value* | *p value* |
| --- | --- | --- | --- | --- |
| **EO** | 3.36 ± 0.27 | 1.53 ± 0.27 | 8.77 | <0.001* |
| **EC** | 1.74 ± 0.12 | 1.41 ± 0.12 | 2.68 | 0.007* |

Supplementary Table S6: Results of post-hoc pairwise contrasts between intervention protocols for baseline-normalized entire delta frequency activity (0.5-3.9 Hz) during resting state are presented below. The table also shows the estimated marginal means and standard errors (mean ± SE) of baseline-normalized 0.5-3.9 Hz power of each stimulation condition.

*Before Sleep*

| Condition | *Estimated Marginal Mean ± SE*  *rTMS+tACS Group* | *Estimated Marginal Mean ± SE*  *Sham Group* | *t value* | *p value* |
| --- | --- | --- | --- | --- |
| **EO** | 0.88 ± 0.14 | 1.04 ± 0.14 | -1.39 | 0.16 |
| **EC** | 0.95 ± 0.06 | 0.95 ± 0.06 | 0.04 | 0.97 |

*After Sleep*

| Condition | *Estimated Marginal Mean ± SE*  *rTMS+tACS Group* | *Estimated Marginal Mean ± SE*  *Sham Group* | *t value* | *p value* |
| --- | --- | --- | --- | --- |
| **EO** | 2.29 ± 0.14 | 1.06 ± 0.14 | 11.02 | <0.001* |
| **EC** | 1.51 ± 0.06 | 1.03 ± 0.06 | 7.40 | <0.001* |

**Supplementary Table S7-8:**

We employed linear mixed models to assess the main effects and interactions of stimulation protocol (2 levels), sleep stages (5 levels), EEG channel (22 levels) as within-subjects factors on baseline-normalized delta power (0.75 Hz and 0.5-3.9 Hz) during sleep.

In case of significant effects of these primary analyses (Table 4), post-hoc pairwise comparisons using the estimated marginal means approach were performed between stimulation protocols for each sleep stage separately.

Asterisks indicate significant differences.

Supplementary Table S7: Results of post-hoc pairwise contrasts between intervention protocols for baseline-normalized 0.75Hz activity during sleep are presented below. The table also shows the estimated marginal means and standard errors (mean ± SE) of baseline-normalized 0.75 Hz power of each stimulation condition.

| Sleep Stage | *Estimated Marginal*  *Mean ± SE*  *rTMS+tACS Group* | *Estimated Marginal Mean ± SE*  *Sham Group* | *t value* | *p value* |
| --- | --- | --- | --- | --- |
| **N1** | 0.77 ± 1.34 | 1.38 ± 1.33 | -0.60 | 0.55 |
| **N2** | 6.55 ± 1.33 | 7.75 ± 1.32 | -1.20 | 0.23 |
| **N3** | 36.18 ± 1.31 | 32.42 ± 1.31 | 3.89 | <0.001* |
| **REM** | 1.31 ± 1.32 | 1.42 ± 1.32 | -0.11 | 0.91 |

Supplementary Table S8: Results for post-hoc pairwise contrasts between intervention protocols for baseline-normalized entire delta frequency activity (0.5-3.9 Hz) during sleep are presented below. The table also shows the estimated marginal means and standard errors (mean ± SE) of baseline-normalized 0.5-3.9 Hz power of each stimulation condition.

| Sleep Stage | *Estimated Marginal Mean ± SE*  *rTMS+tACS Group* | *Estimated Marginal Mean ± SE*  *Sham Group* | *t value* | *p value* |
| --- | --- | --- | --- | --- |
| **N1** | 1.04 ± 0.45 | 1.47 ± 0.45 | -1.29 | 0.20 |
| **N2** | 5.05 ± 0.45 | 5.33 ± 0.44 | -0.90 | 0.37 |
| **N3** | 20.09 ± 0.44 | 18.15 ± 0.45 | 6.25 | <0.001* |
| **REM** | 1.29 ± 0.45 | 1.57 ± 0.45 | -0.91 | 0.37 |

**Supplementary Table S9-10:**

We employed linear mixed models to assess the main effects and interactions of stimulation protocol (2 levels) and timepoint (3 levels) as within-subjects factors on baseline-normalized global efficiency at 0.75 Hz and between 0.5-3.9 Hz.

The table shows the results of the post-hoc pairwise comparisons using the estimated marginal means for the main effect of protocol on global efficiency at 0.75 Hz and between 0.5-3.9 Hz which was significant in the effect analysis (Table 5 and 6) based on linear mixed models.

The analyses were conducted separately for the resting state in eyes open (EO) and eyes closed (EC) conditions.

Supplementary Table S9: Results of post-hoc pairwise contrasts between rTMS and tACS Trough and sham stimulation protocols for baseline-normalized global efficiency at 0.75Hz during the resting state are presented below. The table also shows the estimated marginal means and standard errors (mean ± SE) of baseline-normalized global efficiency at 0.75Hz of each stimulation condition.

*Before Sleep*

| Condition | *Estimated Marginal Mean ± SE*  *rTMS+tACS Group* | *Estimated Marginal Mean ± SE*  *Sham Group* | *t value* | *p value* |
| --- | --- | --- | --- | --- |
| **EO** | 0.87 ± 0.08 | 1.07 ± 0.09 | -1.71 | 0.09 |
| **EC** | 0.98 ± 0.05 | 1.01 ± 0.04 | -0.42 | 0.68 |

*After Sleep*

| Condition | *Estimated Marginal Mean ± SE*  *rTMS+tACS Group* | *Estimated Marginal Mean ± SE*  *Sham Group* | *t value* | *p value* |
| --- | --- | --- | --- | --- |
| **EO** | 1.00 ± 0.08 | 1.31 ± 0.08 | -2.84 | 0.006* |
| **EC** | 1.07 ± 0.05 | 1.13 ± 0.05 | -0.92 | 0.36 |

Supplementary Table S10: Results for post-hoc pairwise contrasts between intervention protocols for baseline-normalized global efficiency across 0.5-3.9 Hz during the resting state are presented below. No post-hoc tests were performed for EC, as no significant main effects of stimulation condition, timepoint, or their interaction were found. The table also shows the estimated marginal means and standard errors (mean ± SE) of baseline-normalized global efficiency across 0.5-3.9 Hz of each stimulation condition.

*Before Sleep*

| Condition | *Estimated Marginal Mean ± SE*  *rTMS+tACS Group* | *Estimated Marginal Mean ± SE*  *Sham Group* | *t value* | *p value* |
| --- | --- | --- | --- | --- |
| **EO** | 0.91 ± 0.03 | 1.04 ± 0.03 | -3.77 | <0.001* |

*After Sleep*

| Condition | *Estimated Marginal Mean ± SE*  *rTMS+tACS Group* | *Estimated Marginal Mean ± SE*  *Sham Group* | *t value* | *p value* |
| --- | --- | --- | --- | --- |
| **EO** | 0.93 ± 0.03 | 0.99 ± 0.03 | -1.84 | 0.07 |

**Supplementary Table S11-12:**

We employed linear mixed models to assess the main effects and interactions of stimulation protocol (2 levels) and sleep stages (5 levels) as within-subjects factors on baseline-normalized global efficiency at 0.75 Hz and between 0.5-3.9 Hz, as shown in Table 7.

Post-hoc pairwise comparisons using the estimated marginal means approach were performed between stimulation protocols for each sleep stage separately. Asterisks indicate significant differences.

Supplementary Table S11: Results of the post-hoc pairwise contrasts between intervention protocols for baseline-normalized global efficiency at 0.75Hz during sleep are presented below. The table also shows the estimated marginal means and standard errors (mean ± SE) of baseline-normalized global efficiency at 0.75Hz of each stimulation condition.

| Sleep Stage | *Estimated Marginal Mean ± SE*  *rTMS+tACS Group* | *Estimated Marginal Mean ± SE*  *Sham Group* | *t value* | *p value* |
| --- | --- | --- | --- | --- |
| **N1** | 0.75 ± 0.07 | 0.77 ± 0.06 | -0.27 | 0.79 |
| **N2** | 0.72 ± 0.06 | 0.48 ± 0.06 | 2.84 | 0.005* |
| **N3** | 0.66 ± 0.06 | 0.59 ± 0.06 | 0.95 | 0.34 |
| **REM** | 0.76 ± 0.06 | 0.74 ± 0.06 | 0.21 | 0.84 |

Supplementary Table S12: Results for post-hoc pairwise contrasts between intervention protocols for baseline-normalized global efficiency between 0.5-3.9 Hz during sleep are presented below. The table also shows the estimated marginal means and standard errors (mean ± SE) of baseline-normalized global efficiency across 0.5-3.9 Hz of each stimulation condition.

| Sleep Stage | *Estimated Marginal Mean ± SE*  *rTMS+tACS Group* | *Estimated Marginal Mean ± SE*  *Sham Group* | *t value* | *p value* |
| --- | --- | --- | --- | --- |
| **N1** | 0.54 ± 0.04 | 0.63 ± 0.04 | -1.56 | 0.12 |
| **N2** | 0.50 ± 0.04 | 0.35 ± 0.04 | 2.57 | 0.01* |
| **N3** | 0.41 ± 0.04 | 0.41 ± 0.04 | 0.01 | 0.99 |
| **REM** | 0.55 ± 0.04 | 0.59 ± 0.04 | 0.81 | 0.42 |

**Supplementary Table S13**

Results for paired t-tests between intervention protocols for fast and slow spindle counts in frontal and parietal channels across N2 and N3 sleep stages are shown below. The tables also show the means and standard errors (mean ± SE) of spindle counts of each stimulation condition.

**Fast Spindles (12-15 Hz)**

| Region of Interest | Sleep Stage | *Mean ± SE*  *rTMS+tACS Group* | *Mean ± SE*  *Sham Group* | *t value* | *p value* |
| --- | --- | --- | --- | --- | --- |
| **Frontal** | **N2** | 128.29 *±* 21.80 | 123.08 *±* 23.04 | 0.37 | 0.95 |
|  | **N3** | 18.39 ± 4.59 | 15.84 *±* 3.26 | 0.70 | 0.95 |
| **Parietal** | **N2** | 176.43 ± 32.41 | 181.70 ± 35.28 | -0.32 | 0.81 |
|  | **N3** | 32.84 ± 7.39 | 29.42 ± 5.82 | 0.55 | 0.81 |

**Slow Spindles (9-11.9 Hz)**

| Region of Interest | Sleep Stage | *Mean ± SE*  *rTMS+tACS Group* | *Mean ± SE*  *Sham Group* | *t value* | *p value* |
| --- | --- | --- | --- | --- | --- |
| **Frontal** | **N2** | 56.7 *±* 7.59 | 54.06 *±* 9.09 | 0.45 | 0.66 |
|  | **N3** | 52.03 ± 22.22 | 68.43 *±* 29.97 | -1.84 | 0.25 |
| **Parietal** | **N2** | 85.29 ± 17.14 | 86.21 ± 14.79 | -0.10 | 0.93 |
|  | **N3** | 28.93 ± 9.09 | 25.50 ± 6.00 | 0.80 | 0.87 |

**Supplementary Table S14**

Results of the paired t-tests between intervention protocols for the ratio of each sleep stage are presented below. The table also shows the means and standard errors (mean ± SE) of the ratio (%) of each sleep stage in each stimulation condition.

| Sleep Stage | *Mean ± SE*  *rTMS+tACS Group* | *Mean ± SE*  *Sham Group* | *t value* | *p value* |
| --- | --- | --- | --- | --- |
| **N1** | 7.44 ± 1.03 | 6.60 ± 1.23 | 0.75 | 0.88 |
| **N2** | 49.67 ± 2.51 | 49.31 ± 2.97 | 0.16 | 0.88 |
| **N3** | 31.01 ± 2.89 | 31.46 ± 3.59 | -0.17 | 0.88 |
| **REM** | 11.88 ± 1.31 | 12.63 ± 1.80 | -0.39 | 0.88 |

**Supplementary Table S15**

Results of the paired t-tests between intervention protocols for sleep onset latency and sleep efficiency are presented below. The table also shows the means and standard errors (mean ± SE) of the sleep onset latency in minutes and of sleep efficiency in percentage for each stimulation condition.

|  | *Mean ± SD*  *rTMS+tACS Group* | *Mean ± SD*  *Sham Group* | *t value* | *p value* |
| --- | --- | --- | --- | --- |
| **Sleep onset latency** | 5.19 ± 2.00 | 3.88 ± 0.93 | 0.85 | 0.41 |
| **Sleep efficiency** | 83.43 ± 2.97 | 87.74 ± 2.12 | -1.79 | 0.09 |
